# Neural Stem Cells Secreting Bispecific T Cell Engager to Induce Selective Anti-Glioma Activity

**DOI:** 10.1101/2020.07.21.188441

**Authors:** Katarzyna C. Pituch, Markella Zanikou, Liliana Ilut, Ting Xiao, Michael Chastkofsky, Madina Sukhanova, Nicola Bertolino, Daniele Procissi, Christina Amidei, Craig M. Horbinski, Karen S. Aboody, Charles D. James, Maciej S. Lesniak, Irina V. Balyasnikova

## Abstract

Glioblastoma (GBM) is the most lethal primary brain tumor in adults. There is no treatment that provides durable relief for the vast majority of GBM patients. In this study, we’ve tested a bispecific antibody comprised of single-chain variable regions (scFvs) against T cell CD3ε and GBM cell interleukin 13 receptor alpha 2 (IL13Rα2). We demonstrate that this BiTE (BiTE^LLON^) engages peripheral and tumor-infiltrating lymphocytes harvested from patient’s tumors, and in so doing exerts anti-GBM activity *ex vivo*. The interaction of BiTE^LLON^ with T cells and engagement of IL13Rα2-expressing GBM cells stimulates T cell proliferation as well as production of pro-inflammatory cytokines INFγ and TNFα. We have modified neural stem cells (NSCs) to produce and secrete the BiTE (NSCs^LLON^). When injected intracranially in mice with brain tumor, NSCs^LLON^ show tropism for tumor, secrete BiTE^LLON^, and remain viable for several days. When injected directly into tumor, NSC^LLON^ provide significant survival benefit to mice bearing IL13Rα2+ GBM. Our results support further investigation and development of this therapeutic for clinical translation.

## Introduction

Routine treatment of newly-diagnosed GBM consists of surgical resection, chemotherapy, and radiation, which results in a median GBM patient survival of less than 2 years, with just 5% of patients surviving beyond 5 years [1]. Blood-brain barrier (BBB) limiting therapeutic access to tumor, an immunosuppressive tumor microenvironment, and tumor molecular heterogeneity present a unique set of challenges for the development of effective GBM therapies [2–9].

The development of treatments for lessening the immunosuppressive effects of GBM represents an active area of preclinical and clinical neuro-oncology research. Many, if not all, approaches being tested involve increasing T cell cytotoxic anti-tumor activity [10–16]. Large numbers of functional cytotoxic tumor-infiltrating lymphocytes (TILs) correlate with increased progression-free survival for GBM patients [17–20]. However, the immunosuppressive milieu of GBM impairs T cell cytolytic function, reducing the effectiveness of T cell based therapies for treating GBM [21–25]. Numerous lymphocyte-directed therapies are being investigated, including the use of bispecific T cell engagers (BiTEs) [16, 26, 27]. BiTEs can be produced and used without need of patient-specific BiTE individualization, and can therefore be considered as “off the shelf” therapeutics [28–31]. The use of BiTEs targeting tumor-associated antigens (TAAs) has been approved by the Food and Drug Administration (FDA) in treating liquid and solid peripheral tumors, and BiTE-associated treatments are currently being evaluated in multiple clinical studies (*e.g*., NCT03792841, NCT04117958, NCT03319940) [32–34].

BiTEs consist of two single-chain variable fragments (scFvs) connected by a flexible linker [35]. One of the scFvs is directed to a TAA and the other to CD3 epsilon (ε) that is expressed on T cells [36]. BiTEs engage TILs and cancer cells in an MHC-independent manner, and are therefore unaffected by MHC downregulation that occurs in GBM cells [35–38]. The specificity of BiTE’s tumor antigen-directed scFv is imperative to harness the full therapeutic potential of the recombinant molecule [39]. BiTE anti-cancer activity requires BiTE binding to malignant and immune cells simultaneously, single arm binding to tumor antigen or to CD3ε is therapeutically ineffective [40, 41].

The short half-life of BiTEs reduces the duration of any BiTE treatment-associated toxicity, if observed at all, but this BiTE characteristic necessitates frequent BiTE infusion into patient circulation in order to maintain therapeutic levels of BiTE [41–43]. Several approaches have been proposed and tested to compensate for the rate of BiTE biologic life in treating peripheral cancers [43–45]. These include sustained production of recombinant protein by subcutaneous injection of mesenchymal stem cells (MSCs) seeded into a synthetic extracellular matrix scaffold, liver translation of BiTE mRNA, and peritumoral delivery of MSCs secreting BiTEs.

Neural stem cells (NSCs) have inherent advantages as cellular carrier and producer of BiTE for extracellular secretion since they are native to the brain. NSCs have demonstrated tropism to brain tumors in several preclinical models. Here, we investigated NSCs as carriers and producers of BiTEs targeting the tumor-associated antigen IL13Rα2, for their anti-tumor activity using *in vitro* and *in vivo* models of GBM. *In vitro*, BiTEs show significant antitumor activity when used in co-cultures that include T cells harvested from patients’ blood and tumor tissue. *In vivo*, NSCs modified for BiTE synthesis migrate to tumor in the brains animal subjects, while functioning as intra- and peritumoral BiTE producers. Following are details of the results from our experiments in characterizing NSCs that produce IL13Rα2-directed BiTEs, and that we interpret as supporting additional safety and efficacy analysis for their potential clinical translation in treating GBM patients.

## RESULTS

### Development of tumor-targeting bispecific T cell engagers

Bispecific T cell engager (BiTE) targeting IL13Rα2 was generated using single-chain variable region (scFv) of mAb47 against the IL13Rα2 that has been previously described by our group and scFv of the mAb OKT3 directed towards the invariable ε chain of CD 3 (CDε) [46–50]. ScFvs were connected using a flexible glycine/serine linker in the following orientation: mAbOKT3VH-mAbOKT3VL-mAb47VL-mAb47VH (Fig. 1A). Short and long linkers (SL and LL, respectively) composed of one or three Gly4Ser repeats were used to construct BiTE^SLON^ and BiTE^LLON^. BiTE^SLOFF^ and BiTE^LLOFF^ control molecules were generated by replacing the complementary determinant region 3 of the mAb47 light chain with the sequence of the mAb MOPC-21, which prevents IL13Rα2 binding (Fig. 1A) [36]. Polyhistidine (6His) tag was added at the C-terminus of BiTE constructs for BiTE purification and detection. Lentiviral vectors (pLVX-IRES-ZsGreen1) encoding cDNA for each BiTE were constructed and corresponding lentiviral particles were used to transduce HEK293T cells for production of BiTE proteins. Recombinant BiTE proteins were purified from culture supernatants using HisPure resin. Purified BiTE integrity was verified by western blotting using anti-His antibodies (Fig. 1B).

**Figure 1.**
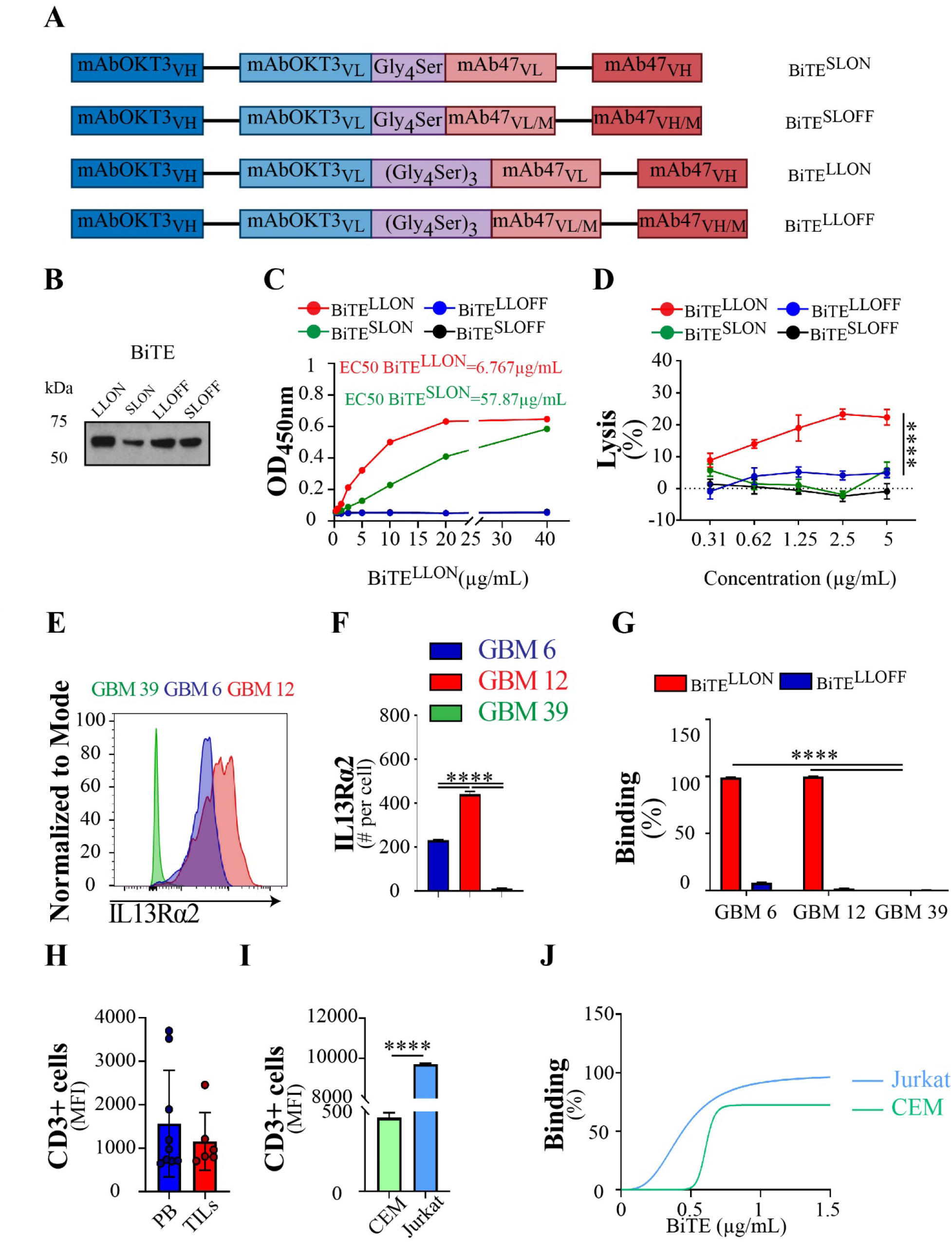
Design of bispecific T cell engager targeting IL13Rα2-expressing gliomas. A) BiTEs consist of the scFv fragments of mAbOKT3 against the CD3ε and scFv of mAb47 against IL13Rα2 connected by either short or long linkers named as BiTE^SLON^ and BiTE^LLON^, respectively. The CDR3 of the light chain (VL/M) of the mA47 was replaced with a sequence of the non-specific MOPC21 antibody to generate BiTE^LLOFF^ and BiTE^SLOFF^ to controls with abridged binding to IL13Rα2. B) Western blotting showed a specific single band after affinity purification and detection with anti-His antibodies at approximately 55 kDa. C) BiTE molecules bound to human IL13Rα2 immobilized on ELISA plates and detected using anti-His tag antibodies. EC50 of BiTE^LLON^ is 6.767 μg/mL, and EC50 of BiTE^SLON^ is 57.87 μg/mL. D) BiTE^LLON^ but not BiTE^LLOFF^ engaged CDε-expressing cells in the killing of IL13Rα2-expressing glioma cells (Two-way ANOVA, n=3-4, ****p<0.0001). E) Example of flow cytometry chromatogram for IL13Rα2-negative GBM39 cell line and IL13Rα2-positive cell lines, GBM6 and GBM12. D) Expression of IL13Rα2 at the cell surface in GBM6, GBM12, GBM39 cell lines (One-way ANOVA, n=3, ****p<0.0001). G) Binding of BiTE^LLON^ and BiTE^LLOFF^ to GBM cell lines (One-way ANOVA, n≥3, ****p<0.0001). H) Expression of CD3ε on the surface of GBM patients’ peripheral blood (PB) lymphocytes and tumor-infiltrating lymphocytes (TILs), (n≥6). I) CD3ε expression on the cell surface of CCRF-CEM, a T lymphoblastoid cell line (CEM) and Jurkat, a T lymphoblast cell line. J) Binding of BiTE^LLON^ to CEM and Jurkat cell lines (n=3).

### Characterization of BiTE binding and function

Binding of purified BiTEs to IL13Rα2 and CD3ε epitopes was examined using ELISA and cell-binding assays. BiTE^LLON^ showed 8.5x higher affinity binding to immobilized IL13Rα2 than BiTE^SLON^ (EC50 values of 6.767 μg/mL and 57.87 μg/mL, respectively: Fig. 1C). Control BiTEs did not bind to IL13Rα2 (Fig. 1C).

All gliomas express IL13Rα2, with GBM expressing the highest levels (SFig. 1A). The ability of IL13Rα2 BiTEs to engage donor T cells in anti-glioma activity was determined using co-cultures of GBM cells with BiTEs and T cells (SFig. 1B, C). BiTE^LLON^ successfully engages T cells, as indicated by BiTE concentration-dependent cytoxicity, with maximal effect observed at 2.5 μg/mL (Fig. 1D, SFig. 1D). BiTE^SLON^ did not induce T cell anti-tumor activity (Fig. 1D).

Patient-derived xenograft cell lines GBM6, GBM12, and GBM39 express different levels of IL13Rα2 (Fig. 1 E, F). BiTE^LLON^ binds to GBM6, and GBM12 cells, but not to GBM39 in which IL13Rα2 is either absent or expressed at a level beneath that required for flow cytometry detection. (Fig. 1G, SFig. 1E). At a concentration of 2μg/mL BiTE^LLON^ saturates IL13Rα2 in GBM6 and 12 (Fig. 1G), and occupies 50% of available receptor in U251 cells (SFig. 1E).

Flow cytometry analysis of GBM patient T cell CD3ε expression revealed substantial interpatient variability in peripheral blood (PB) and tumor infiltrating lymphocyte (TIL) populations. CD3ε-associated fluorescence intensity ranged from 649 to 3058 and from 706 to 1753 for PB and TILs, respectively (Fig. 1 H). We used human lymphocyte Jurkat and CEM cell lines to further test BiTE interaction with CD3 (Fig. 1I, SFig. F, G), and observed saturation binding as BiTE concentrations approached 1.5 μg/mL (Fig. 1J). BiTE CD3ε binding was also apparent when using IL13Rα2 binding defective BiTE^LLOFF^ (SFig. 1H).

To examine BiTE induced T cell anti-tumor activity in vitro we used the chromium51 (Cr51) release assay. BiTE^LLON^ engaged T cell killing of IL13Rα2-expressing GBM6 and GBM12 glioma cells in a T:E ratio-depended fashion (Fig. 2A, B). GBM39 cells, negative for IL13Rα2 expression, were not killed by the T cells, at any tested T: E ratio, in the presence of BiTE^LLON^ and BiTE^LLOFF^ (Fig. 2C). Similar killing activity was observed with T cells harvested from blood of patient with another type of brain tumors, colloidal meningioma, in co-culture with IL13Rα2-positive GBMs in the presence of BiTE^LLON^, but not IL13Rα2-negative cells or BiTE^LLOFF^ (SFig. 2 A-C).At day three of co-culturing, the majority of IL13Rα2-expressing GBM cells were killed by T cells in the presence of BiTE^LLON^, but not in BiTE^LLOFF^ (Fig. 2D). GBM cell killing was associated with proliferation and activation of T cells, as exemplified by the expression of CD69 and CD25 T cell markers. There was no proliferation or expression of activation molecules for T cells co-cultured with BiTE^LLOFF^, or for T cells cultured with BiTE^LLON^ as well as BiTE^LLOFF^ in the absence of IL13Rα2-expressing GBM cells (Fig 2E-G and SFig. 2D-F). T cell proliferation and activation were associated with significant increases in T cell secretion of IL2, IFNγ, and TNFα (Fig. 2 H-J and SFig. G-I).

**Figure 2.**
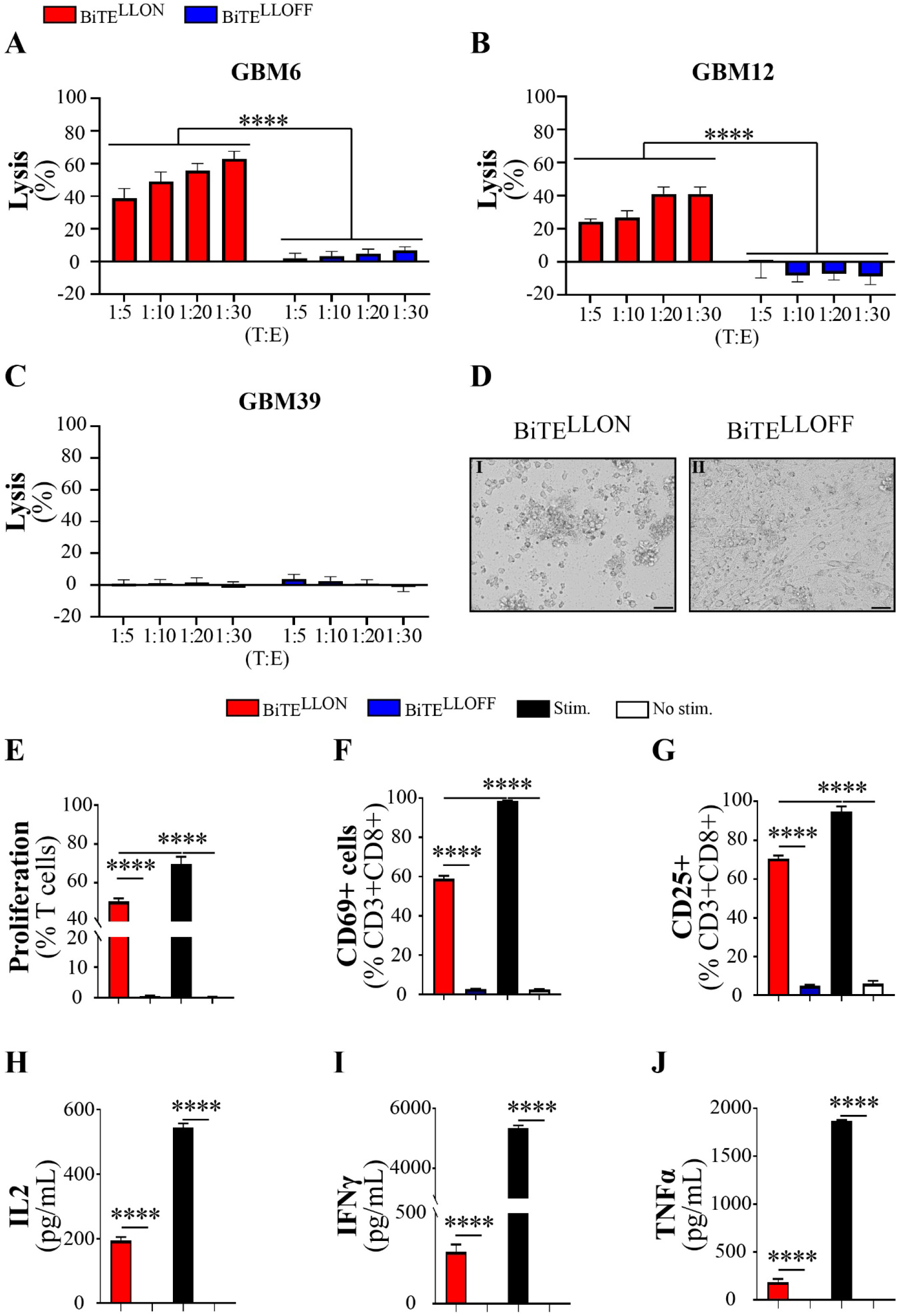
BiTE^LLON^ activates T cell and induces the killing of IL13Rα2+ gliomas. Chromium 51 (^51^Cr) release assay shows that A) GBM6 and B) GBM12 are killed by T cells in the presence of BiTE^LLON^, but not BiTE^LLOFF^ in all tested target-to-effector (E: T) ratios (Two-way ANOVA, n=4, ****p<0.0001). C) GBM39 gliomas are spared by T cells in the presence of either BiTE^LLON^ or BiTE^LLOFF^. D) Bright-field pictures of T cells with IL13Rα2-expressing GBMs after 3 days in co-culture in the presence of BiTE^LLON^(I) and BiTE^LLOFF^(II) (scale bar-50μm). E) Proliferation and expression of activation markers F) CD69 and G) CD25 after co-culture of T cells for 3 days with BiTE^LLON^, BiTE^LLOFF^, stimulation with CD2CD3CD28 beads (Stim.) or absence of any form of stimulation (No stim.) with GBM12 gliomas (One-way ANOVA, n=3, ****p<0.0001). ELISA assay for the production of H) IL2 (24h), I) IFNγ (48h) and J) TNFα (48h) by T cells after co-culture with GBM12 in the presence of BiTE^LLON^, BiTE^LLOFF^, stimulation with CD2CD3CD28 beads (Stim.) or absence of any form of stimulation (No stim.) (One-way ANOVA, n≥4, ****p<0.0001).

### BiTE engages GBM patients’ lymphocytes in an anti-glioma activity

It has been shown that T cells in GBM patients exhibit an exhaustion phenotype characterized by the expression of PD-1/TIM3/Lag3 [7, 51, 52]. Our analysis of patients’ peripheral and tumor-infiltrating lymphocytes indicates cells that are positive for all exhaustion markers (Triple+ cells) comprise 5.48±4.8% of PB PD1+CD8s and 7.67±3.87 of PD1+TILs (Fig. 3A, B). Low level expression of activation markers CD69 and CD2 were detected in CD8+ cells (Fig. 3C, D). In addition, flow cytometry analysis showed that 35.68±5.21% of PB CD8s and 28.99±5.218% of TILs isolated from patients’ tissues are positive for IFNγ (SFig. 3A). The average number of TNFα+ cells within the PB CD8s and TILs was 1.83±0.7% and 8.67±6.17%, respectively (SFig. 3B).

**Figure 3.**
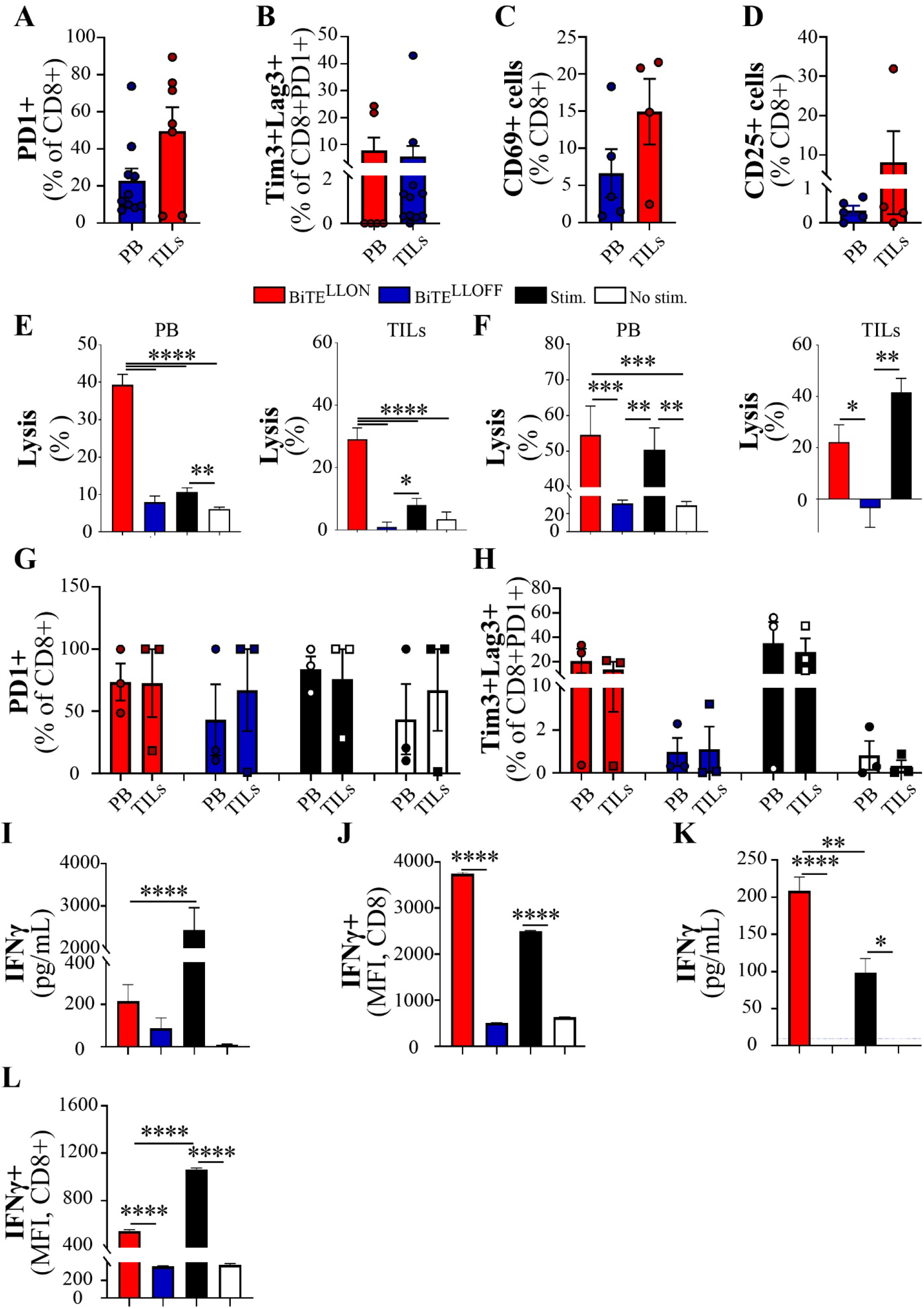
BiTE^LLON^ engages GBM patients’ lymphocytes in an anti-glioma activity. Basal levels of A) PD-1, exhaustion markers B) PD1, Lag3, Tim3 and activation markers C) CD69, D) CD25, in cells isolated from PB and TILs as evaluated by flow cytometry (n≥7). GBM12 ^51^Cr killing assay determined the reengagement of PB and TILs from GBM E) patient1 and F) patient2 into glioma killing in the presence of BiTE^LLON^ and CD2CD3CD28 beads (Stim.), but not BiTE^LLOFF^ or not stimulation (No stim.) (One-way ANOVA, n≥2, *p<0.05, **p<0.01, ***p<0.001, ****p<0.0001). Flow cytometric analysis of G) PD-1 and H) PD-1, Tim3, and Lag3 after co-culture for 3 days of PB lymphocytes or TILs with IL13Rα2-expressing gliomas (n=3). Evaluation of I) secreted (ELISA) and J) intracellular (flow cytometry) levels of IFNγ in lymphocytes isolated from GBM patients’ peripheral blood (One-way ANOVA, n=2-12, *p<0.05, **p<0.01, ****p<0.0001).

We processed blood and tissues from two GBM patients and found that PB and TILs could be stimulated for anti-glioma activity when co-cultured with IL13Rα2+ GBMs in the presence of BiTE^LLON^, but not BiTE^LLOFF^ (Fig. 3E, F). TILs were more potent in killing GBM cells when co-cultured with BiTE^LLON^ as compared to bead-stimulated cells.

PB and TIL co-cultured with GBM cells showed activation and increased expression of PD-1 at the cell surface (Fig. 3G, SFig. 3C,D). There was also an increase of triple+ peripheral lymphocytes and TILs after BiTE^LLON^ and bead stimulation, but not when treated with BiTE^LLOFF^ or in unstimulated conditions (BiTE^LLON^: PB CD8Triple+ 13.52±6.38% and TILs Triple+ 20.52±9.98%; Stimulated: PB CD8 Triple+ 28.03±17.45% and TILs Triple+ 35.15±1.66%, Fig. 3H).

BiTE^LLON^ induced significantly more IFNγ+ PB CD8 cells, as compared to BiTE^LLOFF^ or unstimulated cells, (p<0.0001, Fig. 3I, J). In stratifying cells according to exhaustion markers, we determined that contributors to the production of IFNγ in BiTE^LLON^ stimulated cultures are distributed within all exhaustion marker subtypes, with PD1+Lag3+ cells comprising the largest subgroup (46.53±1.7%, Tab. 1). In positive control PB co-culture (activated with beads), PD-1+ cells were determined as the major producers of IFNγ (Tab. 1). The profile of TNFα in co-cultured PB lymphocytes was similar to the INFγ (SFig. E, F).

Patient TILs secreted significantly more IFNγ when co-cultured with BiTE^LLON^ as compared to negative controls and bead stimulated cells (p<0.005, Fig. 3K). Flow cytometry analysis also showed a higher expression of IFNγ+ in TILs when cultured with BiTE^LLON^ as compared to negative controls, but not to bead activated cultures (Fig. 3L). When stratified by individual exhaustion phenotypes, triple+ TILs were the most significant contributors to IFNγ in BiTE^LLON^ and bead stimulated culture conditions (Tab.1). Flow cytometry analysis revealed that BiTE^LLON^ causes an increase in TIL intracellular TNFα (SFig. 3G). The biggest contributors to the pool of TNFα+ cells were triple+ TILs (Tab.1).

Altogether, these data show that BiTE^LLON^ stimulates anti-glioma activity in patient T cells and induces cytokine production in triple exhausted TILs.

### Development of neural stem cells secreting functional BiTE

BiTEs can cross the BBB, but the rate at which they are degraded and cleared from the body necessitates repeated systemic administration in cancer patients [34, 53]. We and others have demonstrated that neural stem cells (NSCs) can migrate in the brain toward GBM cells/tissue and produce therapeutic antibodies for extended periods of time [54–56]. For this study, we modified immortalized NSCs (NCT03072134) for BiTE^LLON^ and BiTE^LLOFF^ synthesis and secretion (SFig. 4A) [54]. Modified NSCs were FACS sorted by their expression of ZsGreen1 reporter (Fig. 4A). Cytogenic studies of NSCs modified for synthesis and secretion of BiTE^LLON^ (NSC^LLON^) showed the same karyotype as the parental NSCs (SFig. 4B). Results from immunocytochemical analysis (Fig. 4B, C) and western blotting (Fig. 4D) showed that NSC^LLON^ and NSC^LLOFF^ produce BiTE protein. BiTEs secreted by NSC^LLON^, but not NSC^LLOFF^, showed strong binding to hrIL13Rα2 immobilized on ELISA plates (Fig. 4E). Quantitative analysis revealed 1×10^6^ of NSC^LLON^ produced 1.6±0.13 μg of BiTE within the first 24 h. and 2.42±0.26 μg after an additional day in culture (48h) (Fig. 4F). NSC^LLON^ and NSC^LLOFFF^ are tropic for GBM cells in vitro (Fig. 4G, SFig. 4C).

**Figure 4.**
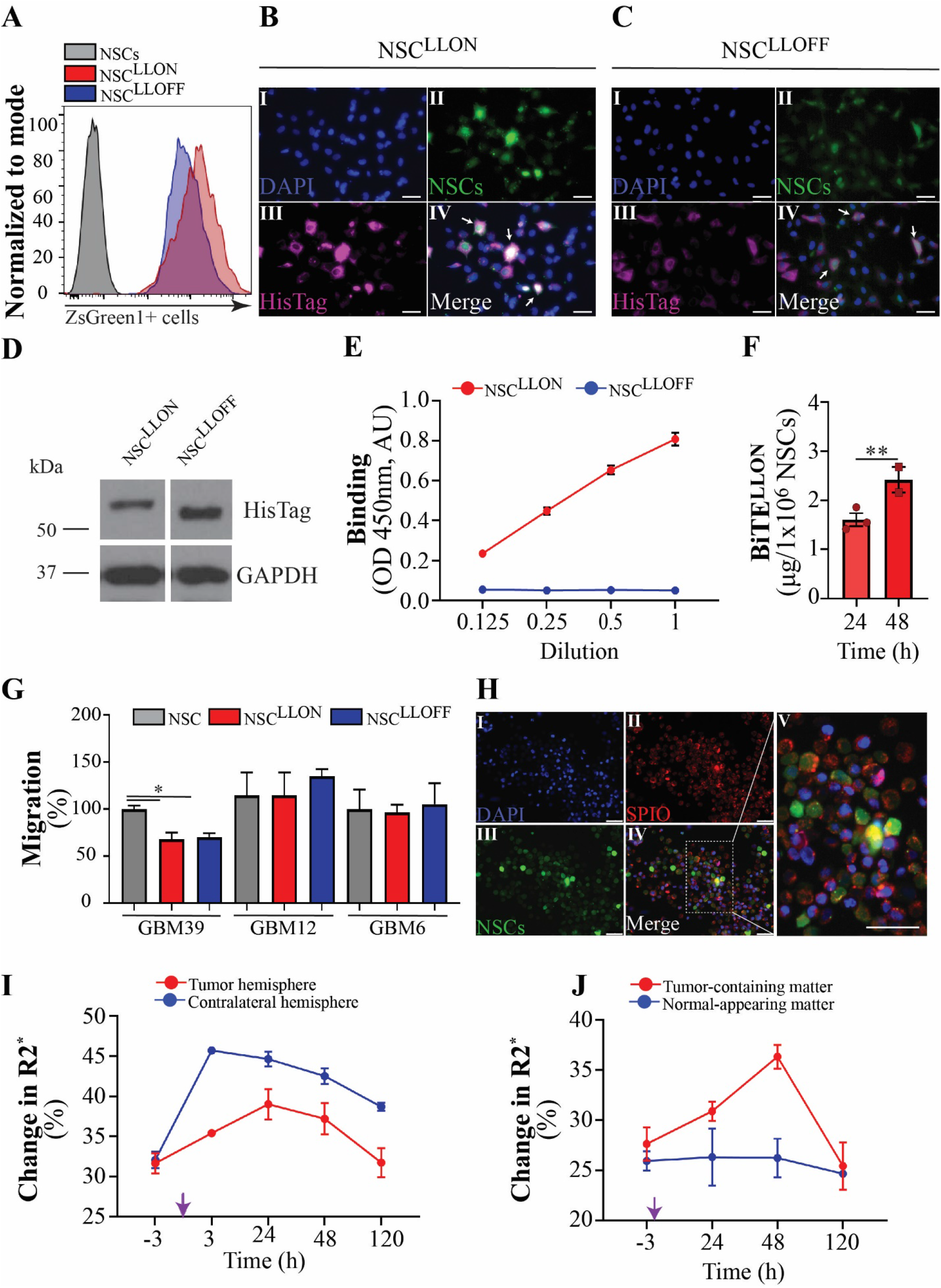
Neural stem cells (NSC) produce and deliver BiTE^LLON^ protein to tumors *in vivo*. BiTE transduced NSCs were sorted for the expression of A) ZsGreen1 (sample histogram) protein and evaluated for the production of BiTE proteins using immunocytochemistry by B) NSC^LLON^ and C) NSC^LLOFF^ cells immobilized on cover glass (cells-DAPI-nucleus(I), NSC-ZsGreen1(II), BiTE protein-α-His-Tag (III), and (IV) merged together, scale bar 50μm), and D) western blotting using anti-His Tag and anti-GAPDH antibodies. E) NSC^LLON^ but not NSC^LLOFF^ produced protein binding to IL13Rα2 protein immobilized on ELISA plates. F) Production dynamics of BiTE^LLON^ by NSC^LLON^ after 24 and 48 hours in culture (T-test, n=3, **p<0.01). Migratory capacity of parental (NSCs) and modified NSC^LLON^ and NSC^LLOFF^ toward human xenograft cell lines: GBM39, GBM12, and GBM6 *in vitro* was assessed in migration assay (One-way ANOVA, n=3, *p<0.05). H) Immunocytochemical analysis NSC^LLON^ labeled with iron particle used for dynamic *in vivo* tracking of NSCs by magnetic resonance imaging (MRI). I) Labeled NSC^LLON^ injected into a contralateral hemisphere, with respect to the tumor, robustly egressed from the injection site and migrated toward tumor (LS Means Pr>|t|=0.0005) and achieved J) maximum tumor coverage within 2 days (LS Means Pr>|t|=0.0063). Purple arrow, injection timepoint=0h.

Iron-labeled NSC^LLON^ were followed by magnetic resonance imaging (MRI) for five days after injection into the hemisphere contralateral to tumor-bearing brain of animal subjects, as previously described [56]. T2* weighted MRI signal showed NSC^LLON^ accumulation in the tumor-bearing hemisphere that reached maximum level at 24h after injection (Fig. 4H, SFig. 4D, E, F). Signal from hemisphere injected with NSC^LLON^ (contralateral to tumor-bearing hemisphere) was highest at 3h after injection, and progressively decreased thereafter (Fig. 4I). Quantitative analysis of R2* data revealed maximal tumor coverage by NSC^LLON^ within 48h from intracranial injection of cells (p=0.0063) (Fig. 4J, and SFig. 4F).

NSCs could be immunosuppressive or immunoreactive due to the presence of major histocompatibility complex I (MHCI) and PDL-1 at their cell surface (SFig. 5A) [57, 58]. NSCs express MHCI on surface (SFig. 5A). Profiling for checkpoint molecules showed that 20.02±2.26% of NSC^LLON^ and 39.57±3.9% NSC^LLOFF^ expressed low level PD-1 ligand (PDL-1) at the cell surface (SFig. 5B,C, D), but this did not affect secreted BiTE^LLON^ from promoting a strong T cell anti-glioma response (NSCs vs NSC supernatant, SFig. 5E). Modified NSCs do not express PDL-2 or cytotoxic T-lymphocyte-associated protein 4 (CTLA4).

NSC^LLON^, but not BiTEs from NSC^LLOFF^, produced BiTE that potently stimulate donor’s lymphocytes leading to killing of IL13Rα2-positive GBM cells (Fig. 5A, B). NSC^LLON^ and NSC^LLOFF^ did not induce lymphocyte killing of IL13Rα2-negative cells (Fig. 5C). GBM patient lymphocytes were also activated against GBM cells by NSC^LLON^ (Fig. 5D). NSC^LLON^, but not NSC^LLOFF^, upregulated lymphocyte expression of granzyme B (GzmB) and Lamp1, markers of degranulation, and mediators of perforin-associated killing of IL13Rα2+ GBM cells. These effects were not observed in co-cultures with IL13Rα2 negative GBM cells (Fig. 5E). BiTEs secreted by the NSC^LLON^ also induced lymphocyte proliferation (Fig. 5F), activation (Fig. 5G, H), and production of type 1 cytokines: IL2, IFNγ, and TNFα (Fig. 5I-K) in co-cultures with IL13Rα2 positive GBM cells (Fig. 5F-K and SFig. 5F-K).

**Figure 5.**
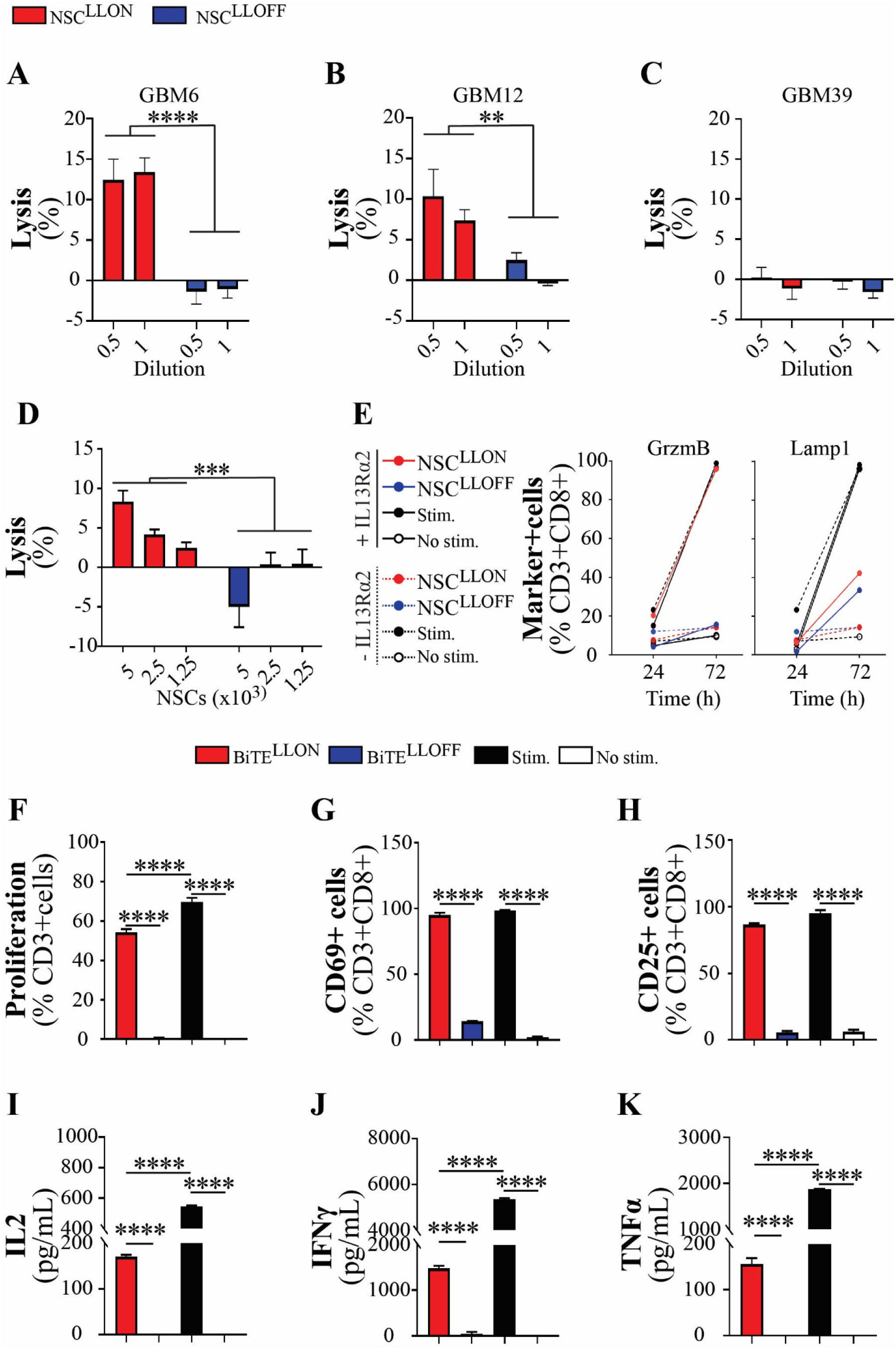
BiTE^LLON^ secreted by NSCs is functional *in vitro*. NSC^LLON^ produce BiTE protein that induced a robust killing in two IL13Rα2+ cell lines, A) GBM6 and B) GBM12 (Two-way ANOVA, n=4, ****p<0.0001). C) BiTE secreting NSCs did not engage donor T cells to kill GBM39, an IL13Rα2-negative cell line. D) NSC^LLON^ presence in a co-culture, but not NSC^LLOFF^, induced reengagement of GBM patient T cells to kill GBM12 cells (Two-way ANOVA, n=3, ***p<0.001). E) Co-culture of donor T cells with GBM12 in the presence of NSC^LLON^ and bead stimulation, but not NSC^LLOFF^ or absence of stimulation, induced expression of Granzyme B (GrzmB) and Lamp1 (markers of cytotoxic T cells activity), F) proliferation and expression of activation markers G) CD69 and H) CD25 in T cells (One-way ANOVA, n≥3, ****p<0.0001). Activation of T cells was associated with the production of I) IL2 (24h), J) IFNγ (48h) and K) TNFα (48h) by T cells after co-culture with GBM12 in the presence of BiTE^LLON^ or stimulation with CD2CD3CD28 beads (Stim.), but not in BiTE^LLOFF^ or absence of any form of stimulation (No stim.) (One-way ANOVA, n≥4, ****p<0.0001).

### NSC^LLON^ administration extends the survival of glioma-bearing animals

We injected NSC^LLON^ proximally (1.5 mm) to GBM12 intracranial tumor in NSG mice (SFig. 6A). Immunohistochemical analysis of sections of brains harvested on days 3 and 7 following NSC^LLON^ injection showed robust infiltration of the tumor mass by stem cells and the production of BiTE protein within the tumor as well as at tumor periphery (Fig. 6A, SFig. 6B,C)). We evaluated NSC^LLON^ for duration of in vivo viability by histopathologic analysis of resected mouse brains from animal subjects that were euthanized at 90 day following NSC^LLON^ injection (SFig. 7A, B), which revealed residual NSC^LLON^ near the third ventricle (SFig. 7B). Moreover, implantation of NSC^LLON^ in tumor-bearing mice in the absence of T cells did not affect the tumor progression as compared to untreated control mice (median: NT-22 days, NSC^LLON^-26 days, Log-rank p=0.269, SFig. 7C).

**Figure 6.**
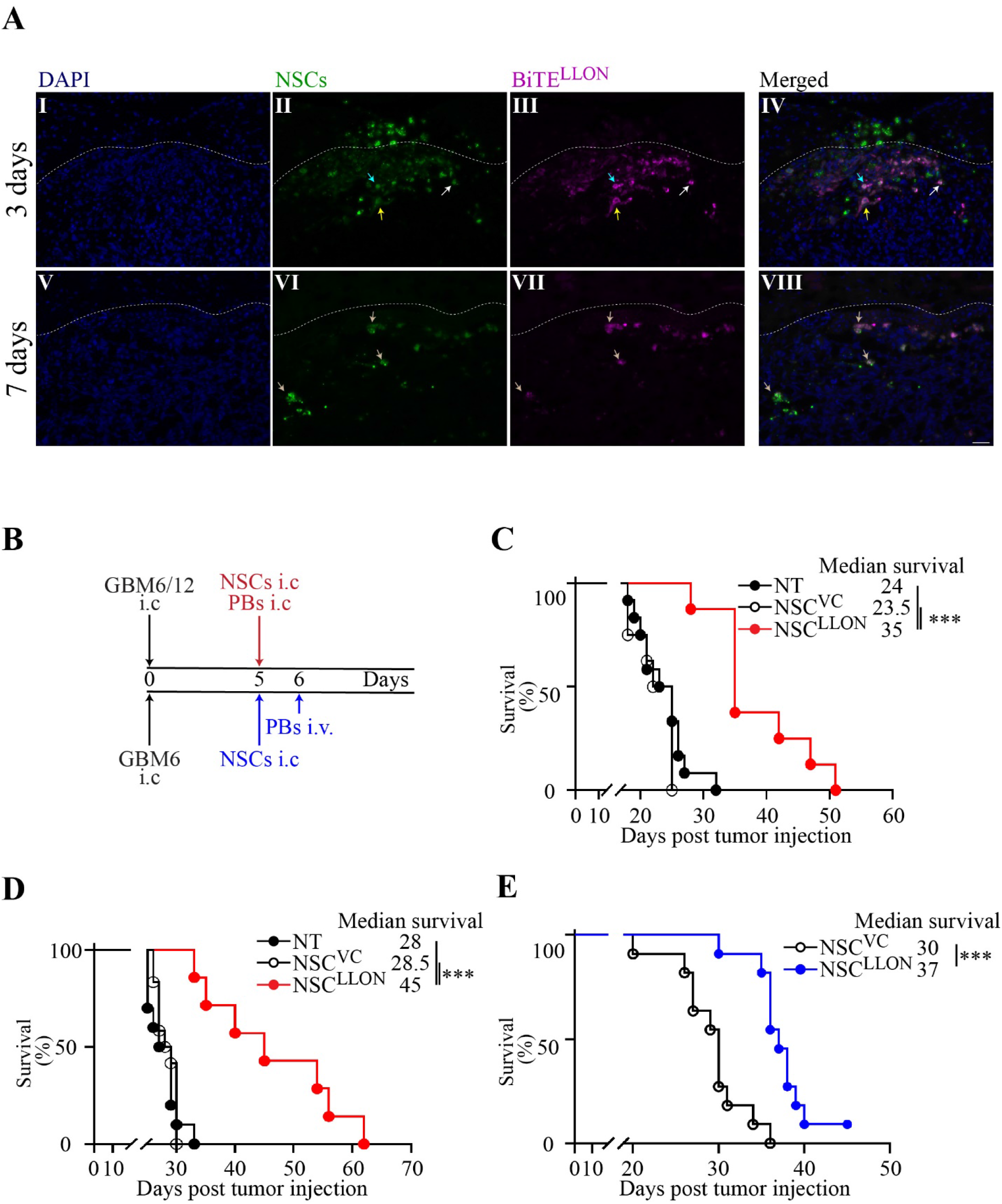
NSC^LLON^ treatment significantly extends the survival of glioma-bearing animals. NSC^LLON^ injected superior and 1.5mm proximal to the tumor mass persist for an extended time, infiltrate tumor mass (arrows), and secret therapeutic proteins(arrows) *in situ* (DAPI-cells, blue; NSCs-green, BiTE^LLON^-magenta; I-IV-3 days and V-VIII-7 days after injection; scale-50μm). B) Animals bearing IL13Rα2-expressing gliomas (GBM6 or GBM12) were treated with single intracranial injection (i.c) of NSC^LLON^ and PB cells, either i.c. or injected intravenously (i.v). C) GBMI2-bearing animals treated with NSC^LLON^ and PB i.c survived significantly longer than not treated (NT) or treated with NSC^VC^ and PB (Kaplan-Meier survival curves were compared using log-rank tests, p<0.0001, and adjusted for p-values using Holm-Sidak method: NT vs. NSC^VC^ p=0.31, NSC^VC^ vs. NSC^LLON^ ***p=0.0003, n=8). D) Similarly, treated mice transplanted with GBM6 cells survived significantly longer from the control NSC^VC^ or NT animals (Kaplan-Meier survival curves were compared using log-rank tests, p=0.0002, and adjusted for p-values using Holm-Sidak method: NT vs. NSC^VC^ p=0.54, NSC^VC^ vs. NSC^LLON^ ***p=0.0009, n≥8). E) GBM6-bearing mice treated with an i.c. NSC^LLON^ and i.v PB lived significantly longer than mice transplanted with control NSC^VC^ and i.v. PB animals.

Infiltration of NSC^LLON^ into the tumor mass with accompanied secretion of BiTE^LLON^ could be detected in tumors 3 days after NSC^LLON^ injection (Fig. 6AII, III, IV, SFig 6B). Both cells and BiTE were still detectable in tumor at day 7 post injection, albeit at lower levels (Fig. 6A II vs. 6A VI, Fig. 6A III vs. 6A VII).

We treated GBM12-bearing mice by intratumoral injection with either NSC^LLON^ or NSC vector control (NSC^VC^) +PB lymphocytes (Fig. 6B). Treatment with NSCs secreting therapeutic BiTEs increased average animal survival by 67% (Fig. 6C). Animals bearing GBM6 tumors were treated using the same protocol as for GBM12-bearing animals (Fig. 6B, D). GBM6-bearing mice treated with NSC^LLON^+PB lived on average 63.3% longer than animals that were not treated (NT), or those that received an injection of NSC^VC^+PB (Fig. 6C). In contrast to GBM6 and GBM12, treatment of mice-bearing IL13Rα2-negative GBM39 tumors with NSC^LLON^ and PB did not extend animals’ survival, as compared to NT group (SFig. 7D). Animals bearing GBM6 tumor and treated with intratumoral injection of NSC^LLON^ and intravenous injection of PB also survived significantly longer as compared to mice treated with control NSC^VC^ (Fig. 6E). Histopathologic analysis of brains from the control and treated animals showed no evidence of therapy-related toxicity, such as demyelination, encephalomyelitis, or neuronal loss.

Altogether, these data show that NSCs secreting BiTE^LLON^ deliver a sustained source of therapeutic protein available for local engagement of CD3+ cells for an anti-glioma activity.

## DISCUSSION

In this study, we report the development and functional analysis of NSCs secreting novel proteins that stimulate patient-derived T cell anti-tumor activity *in vivo* and *in vitro*. We show that IL13Rα2-targeted BiTEs secreted by NSC^LLON^ promote T cell killing of IL13Rα2+ tumor cells by engaging tumor cell antigen target with T cell CD3ε. Genetically-modified NSCs produce and secrete BiTE while infiltrating the tumor mass, which significantly extends animals’ survival.

Demonstration of target specificity is important for limiting undesirable side effects from BiTE treatment of cancer patients [29, 31, 42, 59], and for this reason we have developed BiTE targeting IL13Rα2 [46, 47]. Indeed, we established BiTE^LLON^ binding specificity for PDX cells expressing IL13Rα2, but not IL13Rα2-negative GBM, and T cell CD3ε. Such binding promotes T cell-mediated tumor cell death. Maximal BiTE anti-tumor activity was observed at a low concentration of 2.5 μg/mL and a target to effector cell ratio (1:5), suggesting that relatively low amounts of recombinant protein are needed for achieving anti-tumor efficacy of treatment [41].

GBM promotes lymphocyte exhaustion, which is indicated by lymphocyte expression of PD-1, Tim3, and Lag3 [51, 52]. Though TILs used in our studies expressed variable levels of these checkpoint proteins, BiTE^LLON^ nonetheless activated patient-derived peripheral and tumor-infiltrating T cell killing of IL13Rα2+ GBMs. The lack of tumor-killing in co-cultures with nonbinding BiTE^LLOFF^, or in the absence of IL13Rα2 antigenic target, support the specificity of BiTE^LLON^.

It has been shown that nonspecific activation of T cells by targeted immunotherapeutic can lead to aberrant cytokine production and cytokine release storm (CRS) in treated patients [42, 60]. In our study, donor T cells stimulated directly with BiTE^LLON^, or by BiTE^LLON^ secreted by NSCs, produced INFγ, a key factor in mobilizing the immune system to prevent tumor growth [61], and TNFα, but only in the presence of IL13Rα2-expressing tumor cells. Interestingly, we found that CD8+ T cells expressing exhaustion markers (PD-1+Tim3+Lag3+) are the major contributors to the pool of secreted cytokines when lymphocytes are cultured with BiTE^LLON^ in the presence of IL13Rα2-expressing cells. Notably, BiTE^LLON^ was successful in activating both PB lymphocytes and TIL cells against tumor. This occurred even though T cells were harvested from patients receiving steroid treatment, a known suppressor of T cell activity [62, 63].

Inhibition of T cell function by promoting the aberrant expression of immune checkpoint proteins and MHC-downregulation are key mechanisms by which GBM escapes anti-tumor immune response [37, 51, 52]. Immune-checkpoint inhibitor (ICI) therapy for treating GBM has experienced limited success, in part due to BBB restricting ICI access to intracranial tumor [2, 64–66]. It is plausible that direct administration of ICI with NSC^LLON^ into tumor would ensure therapeutic dissemination within target tissue that, in turn, would allow determination of the potential of this combination therapy for counteracting the effects of lymphocyte exhaustion and reduced MHC expression on tumor cells.

Results from MRI tracking of intracranially administered NSC^LLON^ revealed tropism of therapeutic cells for tumor, which is an important property for achieving activity against residual disease following surgical debulking of the central tumor mass. Thus, NSC^LLON^ administered to the walls of a tumor resection cavity should result in NSC migration toward remaining tumor. In addition to tumor tropism, our results show that NSC^LLON^ persist and secrete BiTE in tumor for several days following administration, so that therapeutic is being continuously delivered within tumor.

The animal subjects used in our xenograft models lack a fully functional immune system, and therefore have an inherent deficiency in recapitulating the tumor microenvironment of a GBM patient. Future studies utilizing immunocompetent hosts are important for revealing BiTE interactions with the host immune system [67]. The use of humanized mouse models, in which mice have been modified for durable production of human immune cells, or mice expressing human CD3 receptor, will also prove informative regarding BiTE-immune system interactions.

In conclusion, the results of our study represent a starting point from which to expand investigation of BiTEs, and specifically IL13Rα2-targeting BiTEs produced by genetically modified NSCs, for treating GBM. The results we have in hand show an exciting potential of this therapy for contributing to improved outcomes for GBM patients.

## MATERIALS AND METHODS

### Cell lines and cell culture

HB1.F3.CD human neural stem cells (NSCs) (provided Dr. K.S. Aboody, City of Hope), HEK 293/17 cells (ATCC, Manassas, VA), U251 GBM cells (kindly provided by Dr. M. Gutova, City of Hope) and GBM 6, 12, 39 (provided by Dr. Charles .D. James, Northwestern University), were maintained in a DMEM (Corning, Manassa, VA) supplemented with 10% fetal bovine serum (Atlanta Biologicals, Flowery Branch, GA), GlutaMax (Gibco, Grand Island, NY), and penicillin/streptomycin (Corning, Manassas, VA) and cultured in a humidified incubator supplied with 5% CO_2_. Human peripheral blood mononuclear cells (PB) were isolated from the whole blood of patients or donor buffy coats (Vitalant, former Life Source, Chicago, IL). Isolated CD3+ cells and tumor-infiltrating lymphocytes (TILs) were maintained in Human T Cells Expansion (STEMCELL Technologies, Vancouver, Canada) medium supplemented with 5% of heat-inactivated human AB serum (Omega Scientific, Tarzana, CA), penicillin (100 U/mL), and streptomycin (100 mg/mL; Corning Carlsbad, CA). Mouse T cells with the addition of mercaptoethanol and patients’ T cells were incubated in the presence of 10ng/mL of hrIL2 (PeproTech, Rocky Hill, NJ). The identity of all cell lines was evaluated using short tandem repeat profiling performed in-house at the Genomic Core Facility, Northwestern University. All cell lines were tested for mycoplasma using MycoGuard Mycoplasma PCR Detection Kit (GeneCopoeia, Rockville, MD) every 3–6 months. Cytogenetic analysis of NSCs was performed as we previously described [55].

### Generation of BiTE molecules

Bispecific T cell engager (BiTE) moiety targeting IL13Rα2 was generated using a single-chain variable region (scFv) of mAb 47. BiTE moiety targeting CD3ε was generated using scFv of the mAb OKT3 antibody [36]. scFvs were connected using a flexible glycine/serine linker in the following orientation: mAbOKT3_VH_-mAbOKT3_VL_-mAb47_VL_-mAb47_VH_ (Fig. 1A). Short linker (SL) and long linker (LL) were generated with one Gly4Ser (5aa) or three repeats of Gly4Ser (15aa) to compose BiTE^LLON^ and BiTE^LLOFF^, respectively. BiTE^SLOFF^ and BiTE^LLOFF^ control molecules were generated by replacing the complementary determinant region 3 (CDR3) of the mAb47 light chain with the sequence of the mAb MOPC-21 [36](PMID:). BiTE constructs were tagged with 6 histidines (6His) at the C-terminus. BiTE cDNAs were codon-optimized for the expression in human cells, synthesized and cloned in pAmp vector by the Thermo Fisher Scientific (Waltham, MA). Alexafluo647 labeling kit was used to label generated BiTEs for the binding studies. Using online ExPASy tools (https://www.expasy.org/tools/), we determined that the SL BiTE (VIRT-125382) molecule is 530 amino acids and LL BiTE (VIRT-131651) is 548 amino acid long, with a respective molecular weight of 56.948 kDa for SL and 58.095 kDa for LL (“Quest Calculate™ IgG Concentration Calculator.” AAT Bioquest, Inc, 17 Jan. 2020, https://www.aatbio.com/tools/calculate-IgG-concentration.).

### Generation of 293T and neural stem cells secreting BiTE protein

The pLVX-IRES-ZsGreen1 plasmid (Takara Bio USA, Inc., Mountain View, CA) was used to generate construct encoding BiTE cDNA. Briefly, the plasmid was cut with EcoRI and BamHI restriction enzymes (NEB, Ipswich, MA), and BiTE cDNA excised from the pAmp vector was directly ligated with T4DNA ligase. DH5α cells (NEB, Ipswich, MA) were transformed and grown overnight on ampicillin LB agar plates. Selected colonies were grown in LB broth in the presence of 100μg/mL of ampicillin. Plasmid DNA was purified using reagents and QIAGEN Plasmid Midi Kits protocol (Germantown, MD). Purified DNA was subjected to Sangers sequencing (GENEWIZ, South Plainfield, NJ). Lentivirus plasmids–encoding for BiTEs cDNA and 4^th^ generation Lenti-X™ Packaging Single Shots (Takara Bio USA, Inc., Mountain View, CA) were used to produce lentiviral particles in HEK 293/17 cells. Cell supernatants were collected at 24 and 48 h after transfection of HEK 293/17 cells and concentrated using a LentiX concentrator (Takara Bio USA, Inc., Mountain View, CA). HEK 293/17 cells or NSC cells were plated in 6 well plates at a density of 10^5^ per well, and next day transduced with viral concentrate in the presence of 4 and 1 μg/mL polybrene, respectively (Sigma, St.Louis, MO), and cultured overnight at 37^0^C/5%CO_2_. The following day media was replaced and transduced cells were expanded. Transduced 293T or NSCs were subjected to fluorescence-activated cell sorting (Robert H. Lurie Comprehensive Cancer Center Flow Cytometry Core Facility (RHLCCC), Chicago, IL) to select for cells expressing a comparable level of ZsGreen1 protein among different BiTE lines. Sorted cells were expanded in culture as specified. Before the collection of the supernatant containing the BiTE proteins, 293T cells at 80% confluence were shifted to 32^0^C and incubated for 3 days in the presence of the proteases inhibitors (Sigma, St. Louis, MO) to induced maximal protein production. NSC secreting BiTE proteins were cultured at 37^0^C/5%CO_2_. In some experiments, cells were passed 48h prior to injection to animals to achieve 90% confluency on the day of surgery. Similarly, cells were prepared for the quantification of NSC^LLON^ BiTE production. In the MRI studies, NSC^LLON^ were incubated with bFGF (20ng/mL) to support cell fitness in serum-free labeling conditions.

### Affinity purification and size exclusion chromatography

Supernatant from the 293T cells secreting BiTE proteins were collected and subjected to affinity purification using cOmpleteTM His_tag purification resin (Sigma-Aldrich, St.Louis, MO). In some studies, BiTE proteins were isolated and processed using size exclusion chromatography at the Recombinant Protein Production Core (rPPC) of Northwestern University. Purified BiTE protein were, dialyzed and stored in 50mM sodium phosphate, 300 mM sodium chloride, pH 7.4. All samples were stored at −80°C until use.

### Isolation of PB T cells and tumor-infiltrating lymphocytes

Peripheral blood mononuclear cells (PBMCs) were isolated from the blood of healthy human donors (LifeSource, Rosemont, IL) or patients treated at the Northwestern University Hospital (Nervous System Tumor Bank) with informed consent under the protocols approved by Institutional Review Board (IRB). In some experiments, PBMCs were used to isolate T cells by CD3-negative selection (EasySep Human T cells Isolation, STEMCELL Technologies, Vancouver, Canada). Tumor-infiltrating lymphocytes (TILs) were isolated from perioperative human patients’ brain tissues (Nervous System Tumor Bank) using EasySep Human CD8 T Cells Isolation Kit (STEMCELL Technologies) following the tissue digestion as previously described [68, 69].

### Functional studies

PB CD3+ and CD8+ TILs were used for the co-culture in functional assays. Two types of controls were used in all experiments except when noted: activated cells co-cultured in the presence of CD3/CD28/CD2 activating bead (STEMCELL Technologies, Vancouver, Canada) and non-activated cells co-cultured in the presence of complete media without beads. IL13Rα2+ cells of hrIL13Rα2 were used for antigenic stimulation except when noted. GBM39, IL13Rα2-negative cells, or GBM-free co-cultures were used as antigen specificity controls as noted per experiment. The co-culture experiments were at the range of target to effector cells (T: E) ratio 1:1-1:30 as specified, where the target is glioma cells, and effectors are T cells. In experiments with BiTEs dose-response concentrations of the recombinant protein were as specified. In experiments with fixed BiTE concentrations, 5μg/mL of BiTE protein was used, and in fixed T: E ratios, the 1:20 was used.

The cytotoxic activity of T cells against glioma cells in the presence of the BiTE proteins or NSCs secreting BiTE proteins was determined using standard ^51^Cr release assay. Depending on the assay, increasing T:E cell ratios or BiTE concentration were used as specified in the result section. Released Cr51 readings were obtained at 18-24 h of the co-culture as previously described [48].

The interleukin 2 (IL2), interferon-gamma (IFNγ) and tumor necrosis factor-alpha (TNFα) production by activated T cells was analyzed using ELISA and flow cytometry. Samples of co-culture supernatant were collected at 24, 48, and 72 h, and stored at −80°C prior to ELISA assay. Manufacturer’s recommendations (R&D Systems, Minneapolis, MN) were followed for data generation and acquisition. Functional flow cytometry was performed as previously described [48] using antibodies specified in flow cytometry section.

### BiTE binding studies and quantification

Enzyme-linked immunosorbent (ELISA) assay was performed to determine the binding of BiTE proteins to human IL13Rα2. ELISA was performed in 96-well plates coated with 1 μg/mL of human recombinant IL13Rα2hFc (R&D Systems, Minneapolis, MN). Following blocking with PBS/2%FBS and washes with TBS-Tween 20 buffer (Boston Bioproducts, Ashland, MA), purified BiTE proteins or NSCs supernatants were incubated for 1h room temperature (RT) at various concentrations or dilutions. Bound BiTE proteins were detected with HRP-conjugated anti-HIS antibodies (Abcam, Cambridge, MA) using 1-Step™ Slow TMB-ELISA (Thermo Scientific, Rockford, IL) and 2N HCl according to manufacturer’s directions. Plates were read at 450nm and 570nm using a BioTek plate reader (BioTek, Winooski, VT), and data were plotted using the Microsoft Excel program prior to analysis.

### Flow cytometry

Fluorescently labeled anti-human antibodies against CD45, CD69, CD3, CD4, CD8, CD25, IFN-γ, TNFα, PD-1, PDL-1, Tim 3, Lag 3, PDL-2, CTLA4 and fixable proliferation and viability dyes were used (BioLegend). Data acquisition for BiTE binding to glioma and immune cells, and immune cells’ phenotypes was done using the BD Fortessa flow cytometer (Becton Dickinson, Franklin Lakes, NJ) at the Robert H. Lurie Comprehensive Cancer Center Flow Cytometry Core Facility and analyzed using FlowJo 10 (X).

The staining of the cells with antibodies was performed according to the assay’s protocol. In all flow cytometry analysis, prior to staining with specific antibodies/BiTEs single-cell isolates viability was determined using trypan blue exclusion and fixable viability dye (APC/Cy7, eBioscience, Carlsbad, CA). Samples were incubated for 30 minutes in complete FACS buffer (PBS supplemented with 2% FBS) and blocked with Human TruStain FcX™ blocking reagent (BioLegend, San Diego, CA) prior to the staining with antibodies or BiTEs.

Next, glioma or CD3+ cells were incubated with an assay-specific concentration of BiTE proteins for 1 h, followed by detection with 1μg/mL of AF647-conjugated secondary anti-HIS antibodies (Cell Signaling, Danvers, MA) for 30 min on ice. Samples were then washed with FACS buffer for 5 min at 500*g*, 4°C. A secondary antibody or unstained cells were used as controls to determine the specific binding to BiTE proteins to glioma and immune cells.

Surface and intracellular proteins were detected using fluorophore-conjugated primary antibodies. Samples were incubated with surface antibodies on ice for 15-30 min and washed with FACS buffer for 5 min at 500*g*, 4°C. In order to determine the expression of intracellular proteins, cells were fixed and permeabilized at RT using Fix/Perm kit (eBioscience, Affymetrix, San Diego, CA), and subsequently stained using the primary fluorophore-conjugated antibodies for 1 h on ice. Cell-surface IL13Rα2 density measurement was done on stained GBM 6, 12, 39 for anti-human IL13Rα2 and isotype control using QuantiBRITE™PE* kit (BD Bioscience, San Jose, CA) per manufacturers specification. The receptor density was assessed using calibration standards provided by the manufacturer.

### Animal tumor studies

SCID-bg (C.B-Igh-1b/GbmsTac-Prkdcscid-Lystbg N7) mice obtained initially from Taconic Biosciences were bred in-house per approved study-specific animal protocol by the Northwestern University Institutional Animal Care and Use Committee (IACUC). Mixed-gender animals used for the evaluation of antitumor activity were between the ages of 7-16 weeks.

Glioblastoma PDX lines, GBM 6, 12, 39, were routinely passaged in mouse flanks and subcultured for 2-3 passages prior implantation into animals’ brains. Prior to surgery, cells were collected and reconstituted in normal (0.9%) saline. We injected 1.5×10^5^ of GBM 6 or GBM 39 cells and 1.0×10^5^ of GBM12 cells per animal in 2.5μL total volume.

Mice were anesthetized with a ketamine/xylazine at 115/17 mg/kg dose, and prepped according to the approved protocol. Glioma cells were injected at 2 mm from bregma, 3 mm right of the midline suture, and at 3.5 mm depth from the dura. PBMCs and/or NSC secreting BiTE proteins were injected at the same location as tumors five days post-GBM 6 or GBM 12, or 10 days post-GBM 39 implantations. In some experiments, tumor-bearing animals were treated intratumorally with NSCs and peripherally with single intravenous injection of 7×10^6^ of PBMCs 24 h after NSCs transplantation. In some experiments, coordinates of NSCs injections were experiment-specific as noted. Vector control NSCs (NSC^VC^) used in the *in vivo* studies only were generated as BiTE secreting cell lines using the carrier vector. All animals were hydrated with 0.5 mL of lactated Ringer’s electrolyte solution post-operation. Control or treatment animals were randomly assigned to housing cages and separated by gender. Survival experiments were repeated at least twice.

### MRI studies

NSC^LLON^ were labeled with a Molday ION Rhodamine B following cell conditions previously established in our laboratory [56, 58, 70]. Labeled NSC^LLON^ were injected into the contralateral hemisphere in respect to the tumor of GBM12-bearing mice. All mice at baseline (pre-treatment with NSCs) were imaged to localize the tumor and obtain the native quantitative MR parameters. Following the injection mice were scanned at 3, 24, 48, and 120 h post-NSC^LLON^ transplantation. Each scanning session involved anesthetization of animals using isoflurane mixed with 100% O_2_ as previously described [56]. The MRI scans were performed on a 7 Tesla Clinscan MRI scanner (Bruker) equipped with the Syngo MR VB15 user interface (Siemens) and 12-cm gradient (strength 290 mT/m and slew rate 1160 T/m/s), and data analysis was performed as previously described [56]. After obtaining a tri-axial set of reference images, a 3D T2* weighted multi-gradient echo sequence (mGRE) was used to obtain high resolution images of the brain (140 μm isotropic) with the following parameters: TR=80 msec and 9 echoes (for extraction of R2* maps) with TE=2.7, 6.83, 11.26, 16, 20.13, 25, 29.13, 33.26, 37.39 msec, flip angle (FA) =10 and 4 averages.

### Statistics and data analysis

All statistical analyses of data were performed using GraphPad 8 (Prism, La Jolla, CA), Microsoft Excel, or SAS 9.4 (SAS Institute Inc., Cary, NC). Significance was defined as *p* less than 0.05 in all statistical tests. Data in two groups were analyzed for statistical significance using the unpaired Mann-Whitney test or an unpaired two-tailed Student’s t-test, as indicated. One- or two-way ANOVA was used for multiple groups followed by a Tukey’s or Dunnett’s multiple comparisons test. During the *in vivo* studies, investigators were blinded to the identity of experimental groups. Animal survival was analyzed after the generation of Kaplan-Meier plots using the log-rank method to determine the *p*-value. The half-maximal effective/binding concentration (EC50) of the tested BiTEs in the binding or killing assays was determined by fitting data into the nonlinear regression using GraphPad8. All *in vitro* studies were repeated at least three times, with multiple experimental replicates as indicated per experiment, unless otherwise stated. All data were presented as average, unless otherwise specified, and error bars in graphs represent standard error of measurement (SEM). Significance in the MRI studies was determined using a mixed model with the difference of least square means.

## Author Contributions

I.V.B. conceive the study. K.C.P. and I.V.B designed experiments. K.C.P., M.Z.,.L.I, M.C., M.S., C.A., N.B., D.P. performed experiments. K.C.P., T.X., C.M.C, N.B, D.P. performed data analysis. K.C.P, M.S., C.D.J., K.S.A, .M.S.L., I.V.B. contributed to the manuscript writing.

## Acknowledgments

This research was supported by the NIH grants R33NS101150 (I.V.B), and R01NS106379 (I.V.B.) and P50CA221747 (M.S.L). The Northwestern Nervous System Tumor Bank is supported by the P50CA221747 SPORE for Translational Approaches to Brain Cancer. Histology services were provided by the Northwestern University Mouse Histology and Phenotyping Laboratory, which is supported by NCI P30-CA060553 awarded to the Robert H Lurie Comprehensive Cancer Center. The authors are also grateful Dolonchampa Maji, Sergii Pshenychnyi, the Recombinant Protein Production Core, and the Small Animal Imaging Facility at Northwestern University, for assistance with this project.

## Conflict of Interest

I.V.B., and M.S.L., have a patent for the use of ScFv47 for IL13Rα2-targeted cancer therapies. K.S.A holds a patent covering the use of Neural Stem Cells for cancer therapy.

## Notes

### Competing Interest Statement

I.V.B., and M.S.L., have a patent for the use of ScFv47 for IL13Ralpha2-targeted cancer therapies. K.S.A has a patent covering the use of Neural Stem Cells for cancer therapy.

