## Supplementary Figures for "Neural Stem Cells Secreting Bispecific T Cell Engager to Induce Selective Anti-Glioma Activity"

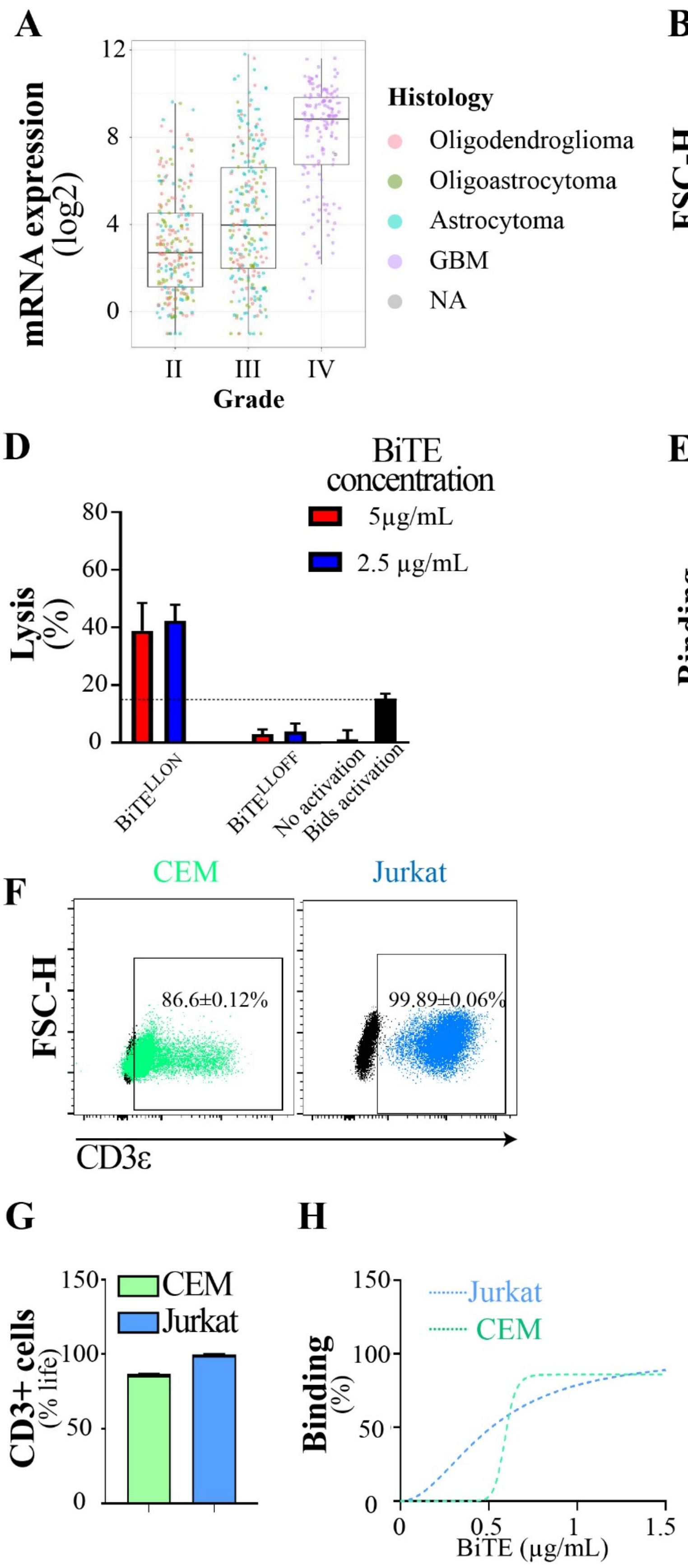

A
mRNA expression (log2)
Grade
II
III
IV
Histology
Oligodendroglioma
Oligoastrocytoma
Astrocytoma
GBM
NA
B
U251
FSC-H
96.13±0.03%
IL13Rα2
C
U251
kDa
75
50
37
IL13α2
β-actin
D
BiTE concentration
5μg/mL
2.5 μg/mL
Lysis (%)
BiTE^LLON
BiTE^LLOFF
No activation
Bids activation
E
U251
EC50=2.276 μg/mL
Binding (%)
BiTE^LLON(μg/mL)
F
CEM
Jurkat
FSC-H
86.6±0.12%
99.89±0.06%
CD3ε
G
CEM
Jurkat
CD3+ cells (% life)
H
Jurkat
CEM
Binding (%)
BiTE (μg/mL)

**Supplemental Figure 1. The activity of bispecific T cell engager targeting IL13Rα2-expressing gliomas**. A) Expression of IL13RA2 at the mRNA level in brain tumors was analyzed using GlioVis data sets (TCGA data set, http://gliovis.bioinfo.cnio.es/). B) Flow cytometry and C) western blotting for the IL13Rα2 in the U251 glioma cell line. D) $BiTE^{LLON}$ engaged CDε -expressing cells isolated from the blood of the patient with meningioma into the killing of IL13Rα2-expressing glioma cells. E) Dose-dependent binding of $BiTE^{LLON}$ to U251 cells. F) CD3ε expression at the cell surface of CCRF-CEM, a T lymphoblastoid cell line (CEM), and Jurkat, a T lymphoblast cell line depicted in a sample dot plot of flow cytometric analysis. G) Quantification of CD3ε in CEM and Jurkat cell lines (n=3). H) Binding of $BiTE^{LLOFF}$ to CEM and Jurkat cell lines (n=3).

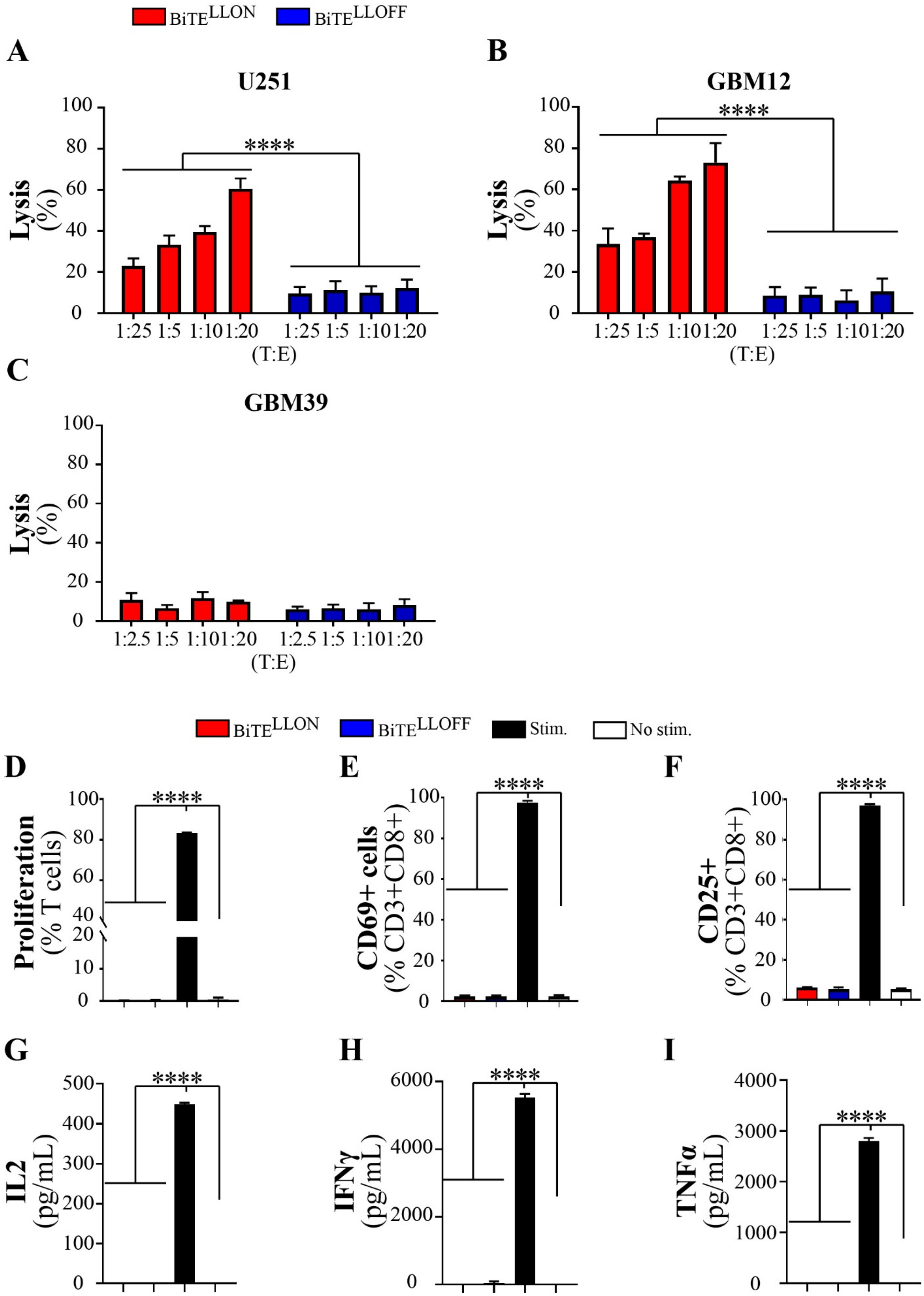

**Supplemental Figure 2. BiTE$^{LLON}$ activates T cell and induces the killing of IL13Rα2+**

**gliomas in an antigen-specific manner**. Chromium 51 (51Cr) release assay shows that A) U251 and B) GBM12 are killed by T cells derived from the blood of a patient with colloidal meningioma in the presence of $BiTE^{LLON}$, but not $BiTE^{LLOFF}$ in all tested target-to-effector (E: T) ratios (Two-way ANOVA, n=3, ****p<0.0001). C) GBM39 gliomas are spared by T cells in the presence of both $BiTE^{LLON}$ and $BiTE^{LLOFF}$. D) Proliferation and expression of activation markers E) CD69 and F) CD25 after co-culture of donor T cells for 3 days with ether $BiTE^{LLON}$, $BiTE^{LLOFF}$, stimulation with CD2CD3CD28 beads (Stim.) or absence of any form of stimulation (No stim.) in the absence of IL13Rα2 (One-way ANOVA, n=3, ****p<0.0001). ELISA assay for the production of G) IL2 (24h), H) IFNγ (48h) and I) TNFα (48h) by donor T cells after co-culture in the presence of ether $BiTE^{LLON}$, $BiTE^{LLOFF}$, stimulation with CD2CD3CD28 beads (Stim.) or absence of any form of stimulation (No stim.) in the absence of IL13Rα2 (One-way ANOVA, n≥4, ****p<0.0001).

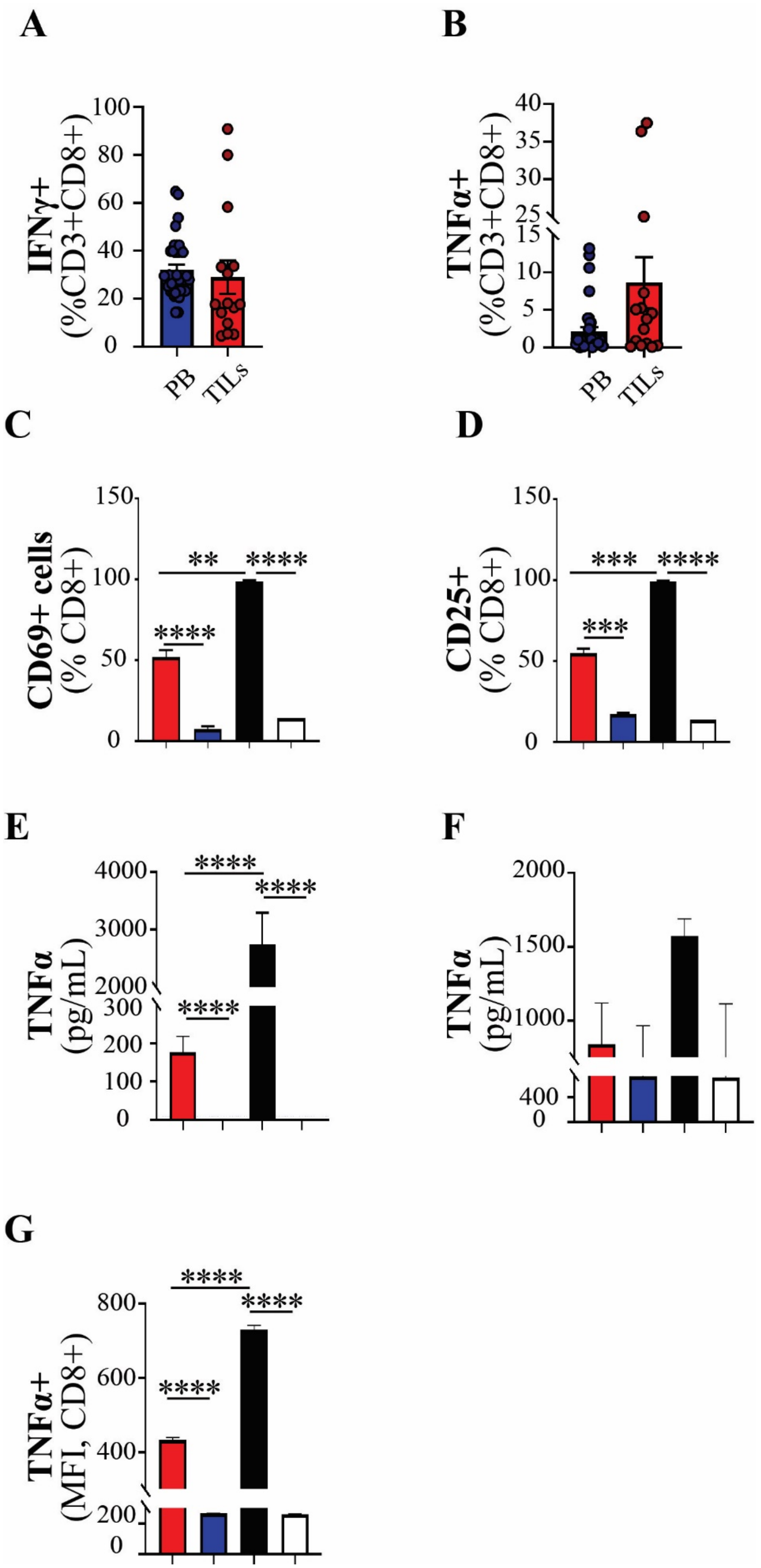

**Supplemental Figure 3. BiTE$^{LLON}$ engages GBM patients' lymphocytes in an anti-glioma**

**activity**. Flow cytometric analysis of basal levels of expression for A) IFNγ, B) TNFα in cells isolated from PB and TILs of glioblastoma patients (n≥7). Flow cytometry analysis for the C) CD69 and D) CD25 markers of T cells activation after co-culture for 3 days with IL13Rα-2 expressing glioma cells in CD3+ lymphocytes isolated from GBM patients' PB (One-way ANOVA, n=2-12, **$p<0.01$, ***$p<0.001$, ****$p<0.0001$). Evaluation of secreted TNFα by ELISA after co-culture for 3 days with IL13Rα-2 expressing glioma cells in E) PB lymphocytes and F) TILs derived from GBM tissue (One-way ANOVA, n=2-12, ****$p<0.0001$). G) Flow cytometric analysis of TNFα expression in TILs harvested from the patients' GBM tissue.

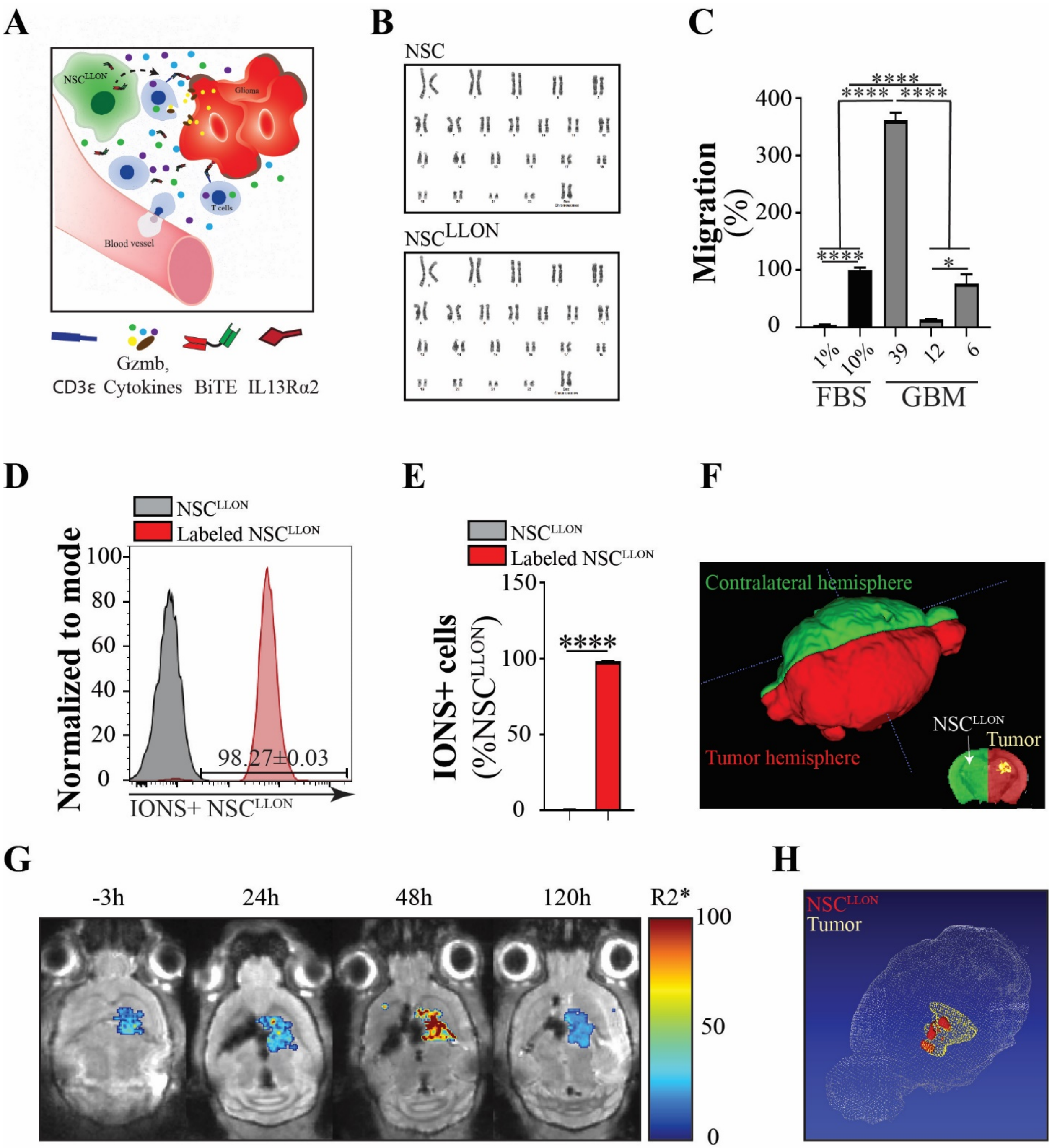

**Supplemental Figure 4. Neural stem cells produce and deliver BiTE$^{LLON}$ protein to tumors.** A) Proposed mechanism of NSC secreting BiTEs activity at the tumor site interacting with 'resident' and newly infiltrating T cells. B) The karyotype analysis of parental NSCs and transduced to produce BiTE$^{LLON}$ (NSC$^{LLON}$). C) Migratory capacity of parental (NSCs) toward human xenograft cell lines: GBM39, GBM12, and GBM6 *in vitro* assessed in migration assay (normalized to the positive control of 10%FBS, One-way ANOVA, n≥3, *p<0.05, ****p<0.0001). D) Histogram and E) the flow cytometry analysis of NSC$^{LLON}$ labeling efficiency with iron particle used for dynamic *in vivo* tracking by magnetic resonance imaging (MRI) (Student T-test, n=3,

****p<0.0001). F) Definition of the contralateral and ipsilateral (hemisphere with the engrafted tumor) hemispheres and anatomical localization of tumor mass used for quantification of MRI data. G) MRI images and pseudo-colored R2* heat-map of labeled $NSC^{LLON}$ migrating into tumor mass, with maximum tumor coverage observed at 2 days (48h). H) 3-D rendition of MRI images representing $NSC^{LLON}$ (in red) tumor coverage at 48h.

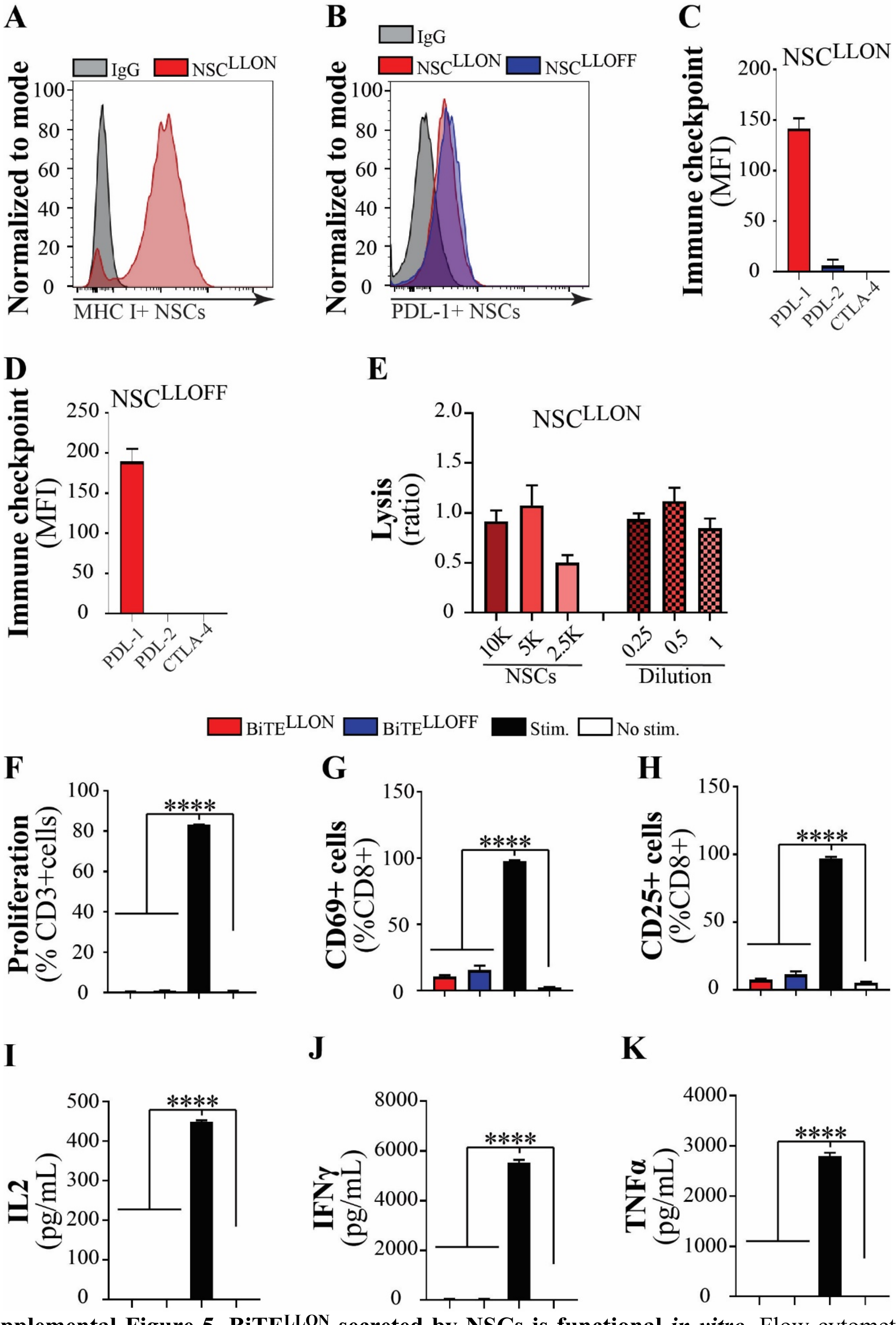

**Supplemental Figure 5. BiTE$^{LLON}$ secreted by NSCs is functional *in vitro*.** Flow cytometric analysis of NSCs secreting BiTEs for the expression of A) MHC I (sample histogram), B) PDL-1 (sample histogram), and quantification of C) PDL-1, PDL-2 and CTLA4 checkpoints (n=3) in $NSC^{LLON}$ and $NSC^{LLOFF}$. E) The presence of BiTE secreted directly by the $NSC^{LLON}$ or contained in supernatants harvested from the $NSC^{LLON}$ successfully engaged GBM patient-derived T cells into a robust killing of U251 cells at the level similar to that of stimulated T cells (ratio=$NSC^{LLON}$/Stimulated T cell responses, n=4). Only bead-stimulated donor T cells, but not lymphocytes stimulated with either $NSC^{LLON}$ or $NSC^{LLOFF}$ or in the absence of stimulation co-cultured without the IL13Rα2 F) proliferated and expressed activation markers G) CD69 and H) CD25 (One-way ANOVA, n≥3, ****p<0.0001). The only activation by beads resulted in the production of I) IL2 (24h), J) IFNγ (48h), and K) TNFα (48h) by T cells after co-culture in the absence of the IL13Rα2 (One-way ANOVA, n≥4, ****p<0.0001).

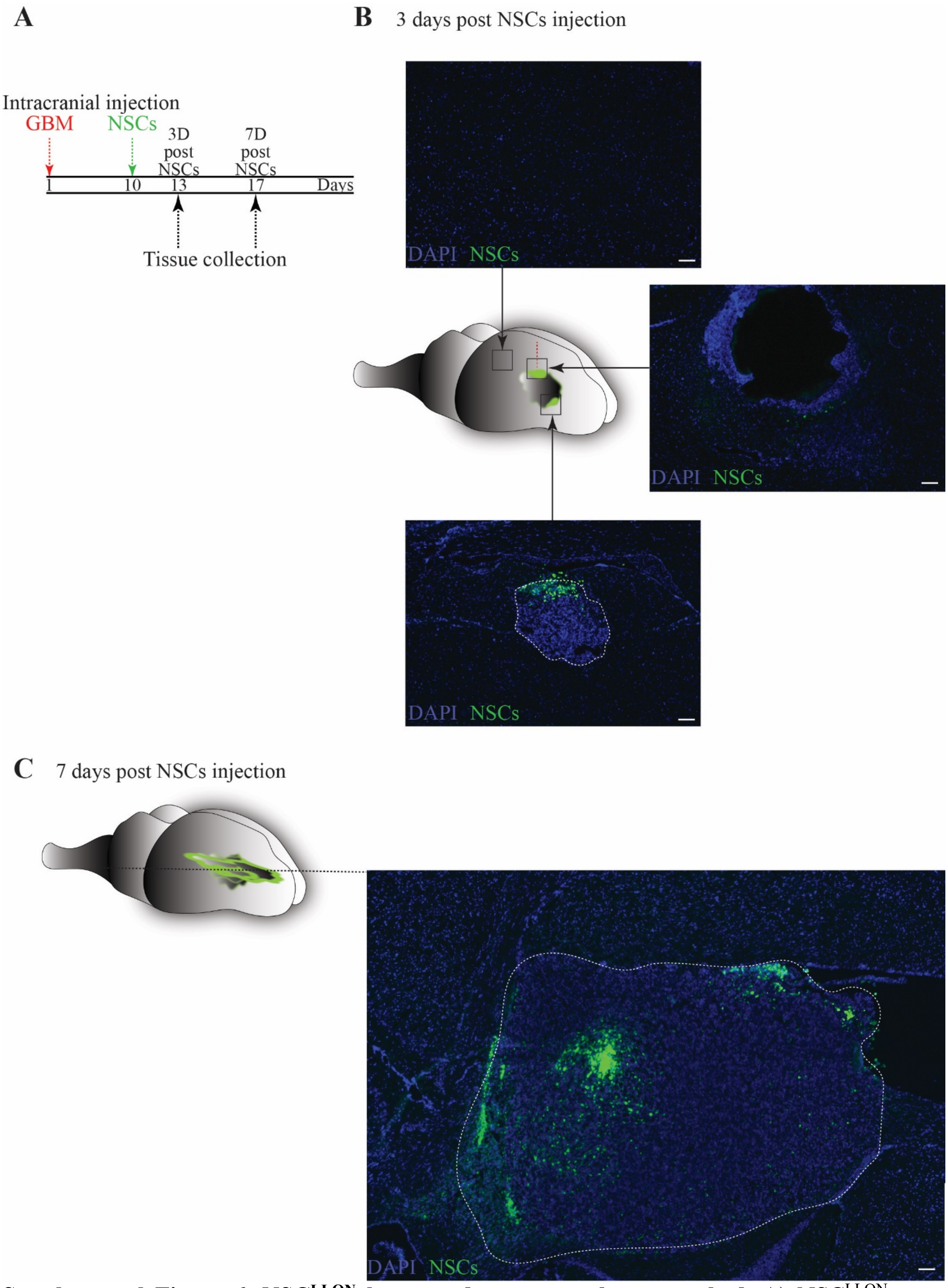

**Supplemental Figure 6. NSC$^{LLON}$ home and penetrate the tumor bed.** A) NSC$^{LLON}$ were injected superior and 1.5mm proximal to established (10 days post-tumor injection) GBM 12 tumor. B) $NSC^{LLON}$ homed into the tumor bed (side and bottom image insets), but not in the normal brain (top image). C) Composite of images obtained at 7 days post-$NSC^{LLON}$ injections through the tumor cross-section shows a robust presence of $NSC^{LLON}$ (DAPI-cells, blue; NSCs-green). Scale-50μm.

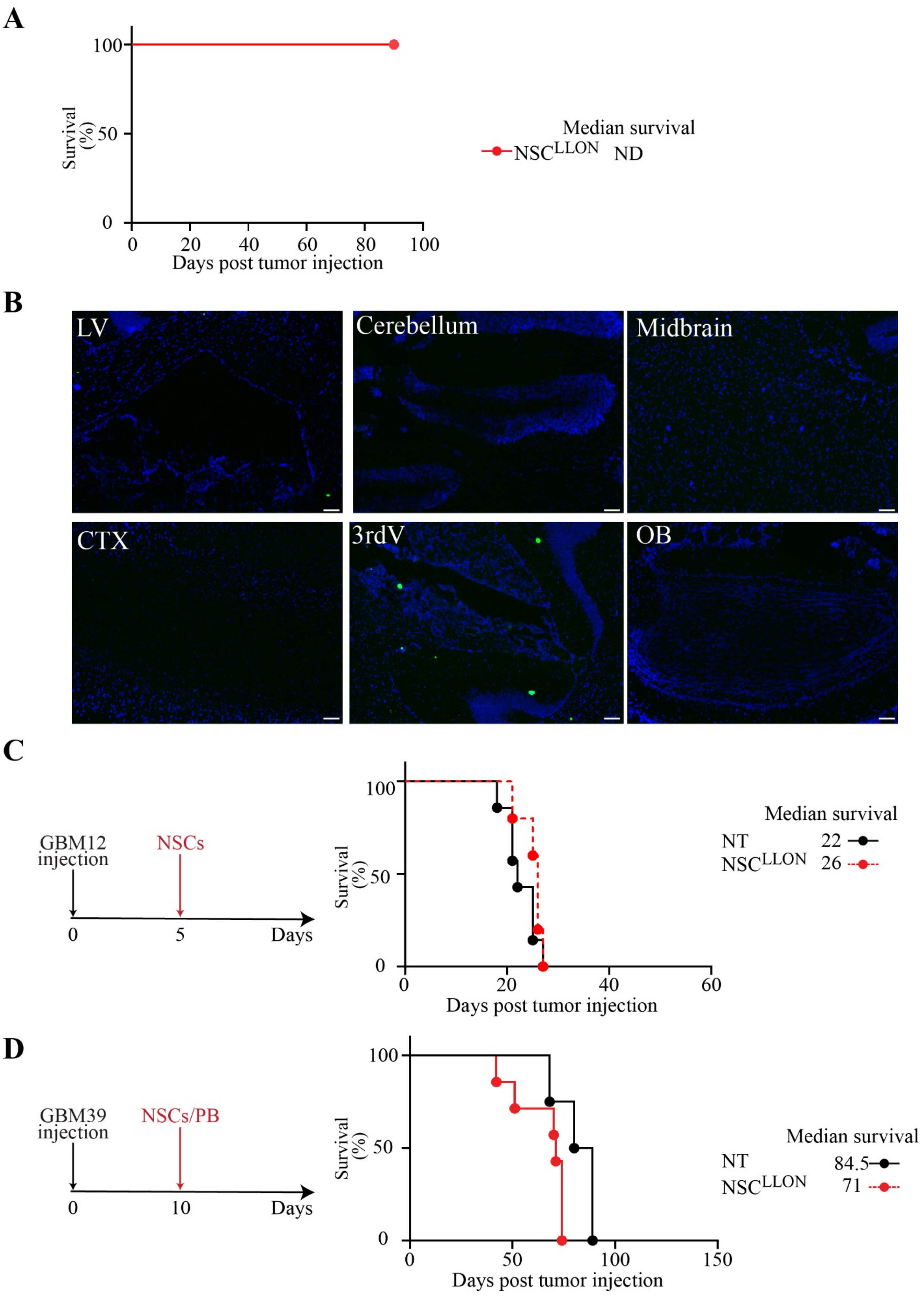

A
100
50
0
Survival (%)
0 20 40 60 80 100
Days post tumor injection
Median survival
NSC^LLON ND
B
LV
Cerebellum
Midbrain
CTX
3rdV
OB
C
GBM12 injection
NSCs
0 5 Days
100
50
0
Survival (%)
0 20 40 60
Days post tumor injection
Median survival
NT 22
NSC^LLON 26
D
GBM39 injection
NSCs/PB
0 10 Days
100
50
0
Survival (%)
0 50 100 150
Days post tumor injection
Median survival
NT 84.5
NSC^LLON 71

**Supplemental Figure 7. NSC$^{LLON}$ activity is antigen-specific and does not support glioma development**. NSC$^{LLON}$ injected into mice (n=3) A) do not develop tumors in mice and B) single cells (as judged by ZsGreen1signal) were observed only in the region of 3$^{rd}$ ventricle, in areas anatomically close to natural NSC niches after 3 months from the intracranial injection (DAPI-cells, blue; NSCs-green). C) NSC$^{LLON}$ do not support GBM12 tumor growth as determined by NSC$^{LLON}$ injection into glioma-bearing mice in the absence of T cells. (Kaplan-Meier survival curve and Log-rank survival analysis, p=27, n= NT-7 and NSC$^{LLON}$-5). D) Treatment with a single intertumoral injection of NSC$^{LLON}$ and PB cells do not extend the survival of mice bearing IL13Rα2-negative GBM39 glioma (n=4-7).

| PB<br>Patients CD3CD8+ cells isolated from peripheral blood | | | | |
|---|---|---|---|---|
| **CD8+PD1+ only cells producing IFNγ** | | | | |
| | BiTE$^{LLON}$ | BiTE$^{LLOFF}$ | Stim. | Not stim. |
| Minimum | 89.7 | 92.4 | 20.2 | 88.6 |
| Maximum | 91.2 | 93.1 | 23.4 | 91 |
| Range | 1.5 | 0.7 | 3.2 | 2.4 |
| Mean | 90.4 | 92.73 | 21.53 | 90.13 |
| Std. Deviation | 0.755 | 0.3512 | 1.665 | 1.332 |
| Std. Error of Mean | 0.4359 | 0.2028 | 0.9615 | 0.7688 |
| **CD8+PD1+Tim3+ cells producing IFNγ** | | | | |
| | BiTE$^{LLON}$ | BiTE$^{LLOFF}$ | Stim. | Not stim. |
| Minimum | 3.53 | 0.92 | 7.04 | 0.11 |
| Maximum | 3.72 | 2.23 | 9.47 | 0.48 |
| Range | 0.19 | 1.31 | 2.43 | 0.37 |
| Mean | 3.607 | 1.7 | 8.52 | 0.2333 |
| Std. Deviation | 0.1002 | 0.6899 | 1.299 | 0.2136 |
| Std. Error of Mean | 0.05783 | 0.3983 | 0.7499 | 0.1233 |
| **CD8+Triple+ cells producing IFNγ** | | | | |
| | BiTE$^{LLON}$ | BiTE$^{LLOFF}$ | Stim. | Not stim. |
| Minimum | 0.46 | 0 | 22 | 0.11 |
| Maximum | 1.17 | 0.15 | 25.1 | 0.12 |
| Range | 0.71 | 0.15 | 3.1 | 0.01 |
| Mean | 0.8133 | 0.1 | 23.4 | 0.1133 |
| Std. Deviation | 0.355 | 0.0866 | 1.572 | 0.005774 |
| Std. Error of Mean | 0.205 | 0.05 | 0.9074 | 0.003333 |
| **CD8+PD1+Lag3+ cells producing IFNγ** | | | | |
| | BiTE$^{LLON}$ | BiTE$^{LLOFF}$ | Stim. | Not stim. |
| Minimum | 4.79 | 4.9 | 44.4 | 8.35 |
| Maximum | 5.94 | 6.57 | 49.9 | 11.1 |
| Range | 1.15 | 1.67 | 5.5 | 2.75 |
| Mean | 5.177 | 5.473 | 46.53 | 9.467 |
| Std. Deviation | 0.6611 | 0.9501 | 2.95 | 1.446 |
| Std. Error of Mean | 0.3817 | 0.5485 | 1.703 | 0.8348 |

| TILs<br>Patients CD8+ cells isolated from tumor tissue | | | | |
|---|---|---|---|---|
| **CD8+PD1+ only cells producing IFNγ** | | | | |
| | BiTE$^{LLON}$ | BiTE$^{LLOFF}$ | Stim. | Not stim. |
| Minimum | 11.6 | 75.3 | 6.14 | 75.8 |
| Maximum | 15.8 | 83.6 | 6.93 | 85.2 |
| Range | 4.2 | 8.3 | 0.79 | 9.4 |
| Mean | 13.97 | 80.47 | 6.53 | 81.37 |
| Std. Deviation | 2.15 | 4.508 | 0.3951 | 4.934 |
| Std. Error of Mean | 1.241 | 2.603 | 0.2281 | 2.849 |
| **CD8+PD1+Tim3+ cells producing IFNγ** | | | | |
| | BiTE$^{LLON}$ | BiTE$^{LLOFF}$ | Stim. | Not stim. |
| Minimum | 12.8 | 3.37 | 5.09 | 4 |
| Maximum | 21.3 | 5.99 | 6.14 | 9.32 |
| Range | 8.5 | 2.62 | 1.05 | 5.32 |
| Mean | 17.9 | 4.937 | 5.587 | 5.987 |
| Std. Deviation | 4.498 | 1.383 | 0.5273 | 2.904 |
| Std. Error of Mean | 2.597 | 0.7987 | 0.3044 | 1.677 |
| **CD8+Triple+ cells producing IFNγ** | | | | |
| | BiTE$^{LLON}$ | BiTE$^{LLOFF}$ | Stim. | Not stim. |
| Minimum | 54.2 | 3.64 | 64.3 | 0.89 |
| Maximum | 59.8 | 6.74 | 69.6 | 7.45 |
| Range | 5.6 | 3.1 | 5.3 | 6.56 |
| Mean | 57.3 | 5.303 | 66.7 | 3.343 |
| Std. Deviation | 2.848 | 1.562 | 2.685 | 3.579 |
| Std. Error of Mean | 1.644 | 0.902 | 1.55 | 2.066 |
| **CD8+PD1+Lag3+ cells producing IFNγ** | | | | |
| | BiTE$^{LLON}$ | BiTE$^{LLOFF}$ | Stim. | Not stim. |
| Minimum | 9.14 | 5.99 | 19.2 | 7.45 |
| Maximum | 11.7 | 14.6 | 22.7 | 12 |
| Range | 2.56 | 8.61 | 3.5 | 4.55 |
| Mean | 10.85 | 9.287 | 21.2 | 9.297 |
| Std. Deviation | 1.478 | 4.646 | 1.803 | 2.393 |
| Std. Error of Mean | 0.8533 | 2.682 | 1.041 | 1.382 |

**Supplementary Table 1. Stratification of INFγ production by exhaustion type in PB lymphocytes and TILs culture.**
